# Human-like Modulation Sensitivity Emerging through Optimization to Natural Sound Recognition

**DOI:** 10.1101/2022.09.25.509427

**Authors:** Takuya Koumura, Hiroki Terashima, Shigeto Furukawa

## Abstract

Natural sounds contain rich patterns of amplitude modulation (AM), which is one of the essential sound dimensions for hearing perception. The sensitivity of human hearing to AM measured by psychophysics takes diverse forms, from low-pass to high-pass, depending on the experimental conditions. Here, we address with a single framework the questions of why such patterns of AM sensitivity have emerged in the human auditory system and how they are realized by our neural mechanisms. Assuming that optimization for natural sound recognition has taken place during human evolution and development, we examined its effect on the formation of AM sensitivity by optimizing a computational model, specifically, a multi-layer (or deep) neural network, for natural sound recognition and simulating psychophysical experiments in which the model’s AM sensitivity was measured. Relatively higher layers in the optimized model exhibited qualitatively and quantitatively similar AM sensitivity to that of humans, even though the model was not designed to reproduce human-like AM sensitivity. The similarity of the model’s AM sensitivity to humans’ correlated with its sound recognition accuracy. Optimization of the model to degraded sounds revealed the necessity of natural AM patterns for the emergence of human-like AM sensitivity. Consistent results were observed from optimizations to two different types of natural sound. Moreover, simulated neurophysiological experiments on the same model revealed a correspondence between the model layers and the auditory brain regions that is based on the similarity of their neural AM tunings. The layers in which human-like psychophysical AM sensitivity emerged exhibited substantial neurophysiological similarity with the auditory midbrain and higher regions. These results suggest that the behavioral AM sensitivity of human hearing has emerged as a result of optimization for natural-sound recognition in the course of our evolution and/or development and that it is based on a stimulus representation encoded in the neural firing rates in the auditory midbrain and higher regions.

## Introduction

Amplitude modulation (AM) is a critical sound feature for hearing (Figure 1). Not only is AM strongly associated with basic hearing sensations such as loudness fluctuation, pitch, and roughness, (Joris et al., 2004), but it is also an essential clue for recognizing natural sounds, including everyday sounds, animal vocalizations, and speech (Dudley, 1939; Shannon et al., 1995; Gygi et al., 2004). Amplitude modulation of a sound is often expressed as a modulation spectrum, i.e., the magnitude of the amplitude change (AM depth) at each modulation-frequency component (AM rate) (Figure 1b, Figure 1c). The significance of AM sensitivity to our hearing functions is supported by its relationship with speech recognition performance. In particular, experiments, mostly conducted on hearing-aid and cochlear implant users, have shown correlations between speech recognition performance and AM detection sensitivity (Fu, 2002; Luo et al., 2008; Won et al., 2011; De Ruiter et al., 2015), the difference in detection sensitivity between low and high AM rates (Cazals et al., 1994), and spectro-temporal modulation sensitivity (Bernstein et al., 2016).

**Figure 1.**
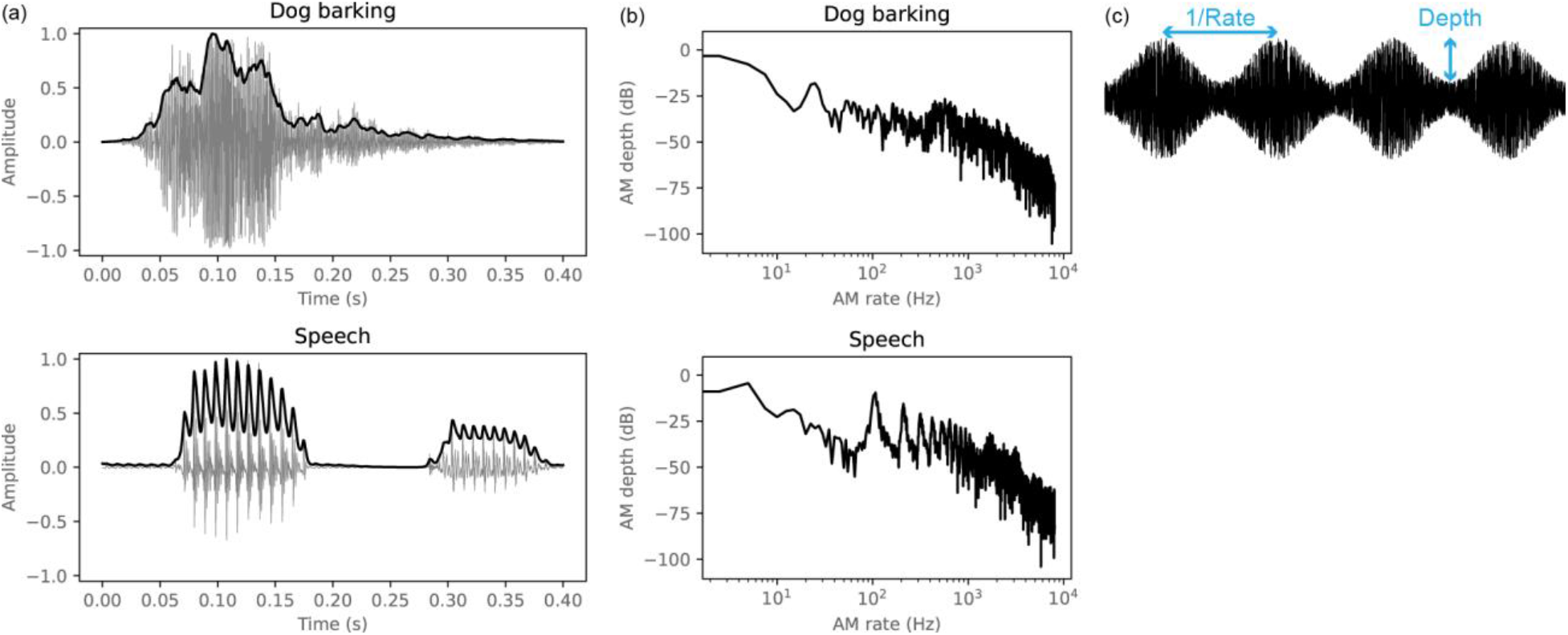
(a) Examples of AM in natural sounds. Excerpts of a dog-barking/speech are shown at the top/bottom. Sound waveforms and their amplitude envelopes are shown by gray and black lines, respectively. (b) Modulation spectra of the sounds in (a). Each sound has a distinct modulation pattern. (c) Illustration of the AM depth and rate (actually, the inverse of the rate) of sinusoidally amplitude-modulated white noise. Generally, the shallower the AM depth is, the more difficult AM becomes to detect.

The properties of AM sensitivity have been investigated mainly through two separate approaches: psychophysics and neurophysiology. On the one hand, psychophysical studies have identified a wide variety of sensitivity curves in the form of the temporal modulation transfer function (TMTF). For humans, TMTF is defined as the AM-detection threshold (i.e., the minimum AM depth required for detection) as a function of AM rate (Figure 2). Typically, it is measured with a sinusoidal AM (Figure 1c). TMTFs measured with a broadband noise carrier have low-pass characteristics (Figure 2, right-most panel) (Viemeister, 1979). The threshold is lower for low AM rates and increases monotonically with increasing AM rate. Studies using AM on narrow-band noise carriers have shown different forms of TMTF (Figure 2, left four panels) (Dau et al., 1997a; Lorenzi et al., 2001a). In particular, the TMTFs show apparent interactions with the carrier bandwidth. The detection thresholds are higher at an AM rate equal to the carrier bandwidth. These carrier-bandwidth-dependent TMTFs have been interpreted in terms of frequency masking in the modulation domain, with the target modulation being masked by the amplitude fluctuation intrinsic to the stimulus mostly due to the narrow carrier bandwidth. Stimulus parameters other than the carrier bandwidth are also different among experiments. They may be additional factors determining the form of the TMTF.

**Figure 2.**
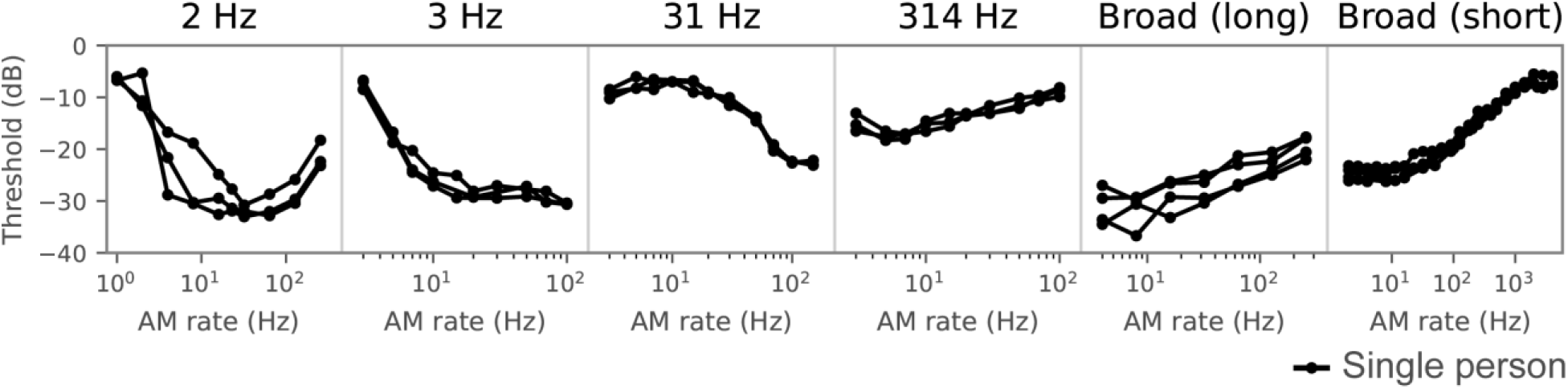
TMTFs of humans, sorted by the carrier bandwidth of the stimulus. The TMTF is defined as the AM detection threshold as a function of the AM rate. Amplitude modulation of broadband carriers yields low-pass-shaped TMTFs with lower thresholds at low AM rates and higher thresholds at high AM rates, whereas it yields high-pass-shaped TMTFs for narrowband carriers. Other stimulus parameters also appear to affect TMTFs. The depicted TMTFs were taken from psychophysics papers (Viemeister, 1979; Dau et al., 1997a; Lorenzi et al., 2001b, 2001a). Each line shows a TMTF in a single person.

On the other hand, neurophysiological studies have found that many neurons throughout the mammalian auditory nervous system (ANS) show tuning to AM (Joris et al., 2004). Their spike rate and/or spike timing depends on the AM rate in the stimulus. Their preferred AM rate varies widely over the range of behaviorally detectable values. There is a systematic change in neuronal tuning at a single-unit level along the auditory pathway where central regions are more sensitive to slow changes and non-linear features (Joris et al., 2004; Sharpee et al., 2011). Although these findings suggest that AM-tuned neurons are somehow involved in behavioral AM sensitivity, the lack of single-unit neural data in humans has made it difficult to establish a direct link with human behavior.

Inspired by the psychophysically observed apparent frequency masking in the modulation domain and neurophysiologically observed AM-tuned neurons, Dau et al. (1997) have proposed that a bandpass filterbank in the modulation domain, called a modulation filter bank (MFB), is involved in auditory signal processing (Dau et al., 1997a). They have built a computational model of auditory signal processing that includes an MFB, with which they have reproduced a variety of psychoacoustic properties including stimulus-parameter-dependent TMTFs (Dau et al., 1997a, 1997b; Derleth et al., 2001; Jepsen et al., 2008). To reproduce a wider range of psychoacoustic phenomena, they have gradually incremented and carefully refined the model components that each performs a specific signal processing computation. Building a model in such a bottom-up fashion is advantageous for describing in detail the effects of different signal processing stages and theorizing on what kinds of signal processing are implemented in the human auditory system. However, to fully understand the properties of AM sensitivity, we should also answer two critical questions: Why has it emerged during our evolution and development? How it is realized by our neural mechanisms?

To provide answers to the “why” and “how” of auditory AM sensitivity, we built a computational model that performs natural sound recognition and compared its psychophysical and neurophysiological properties with those of the human auditory system (Figure 3). First, to investigate why AM sensitivity has emerged, we optimized an artificial deep neural network (NN) for natural sound recognition (Figure 3a) as a way of simulating the optimization that is presumably happening in the auditory system during its evolution and development. We assumed that better recognition of natural sounds yields better evolutionary fitness and hypothesized that natural sound recognition plays a major role in shaping human AM sensitivity (Fu, 2002; Luo et al., 2008; Won et al., 2011; De Ruiter et al., 2015). Then, we froze the learned parameters and simulated psychophysical AM detection experiments on the optimized NN to see whether or not human-like AM sensitivity emerges in some of its layers (Figure 3b). Note that we did not directly model or try to reproduce human AM sensitivity in the optimization process. This kind of two-step optimization-and-analysis procedure has explained a number of properties of the auditory system, such as cochlear frequency tuning (Lewicki, 2002), AM tuning (Khatami and Escabí, 2020), the receptive field in the auditory cortex (Terashima and Okada, 2012), pitch perception (Saddler et al., 2021), sound localization (Francl and McDermott, 2022), and speech processing (Ashihara et al., 2021), as well as in other sensory modalities (Kriegeskorte and Douglas, 2018). To reduce the number of hard-coded assumptions and clarify the relationship between the optimization procedure and the emergent properties, we applied an NN directly to a “raw” sound waveform without any preprocessing (Hoshen et al., 2015; Tokozume and Harada, 2017). This is in contrast with typical auditory models that attempt to implement a hard-coded frequency-decomposition stage in the cochlea (Bruce et al., 2018; Verhulst et al., 2018). We further examined the necessity of the AM patterns inherent in natural sounds and compared our AM detection procedure with the previously proposed temporal-template-based AM detection.

**Figure 3.**
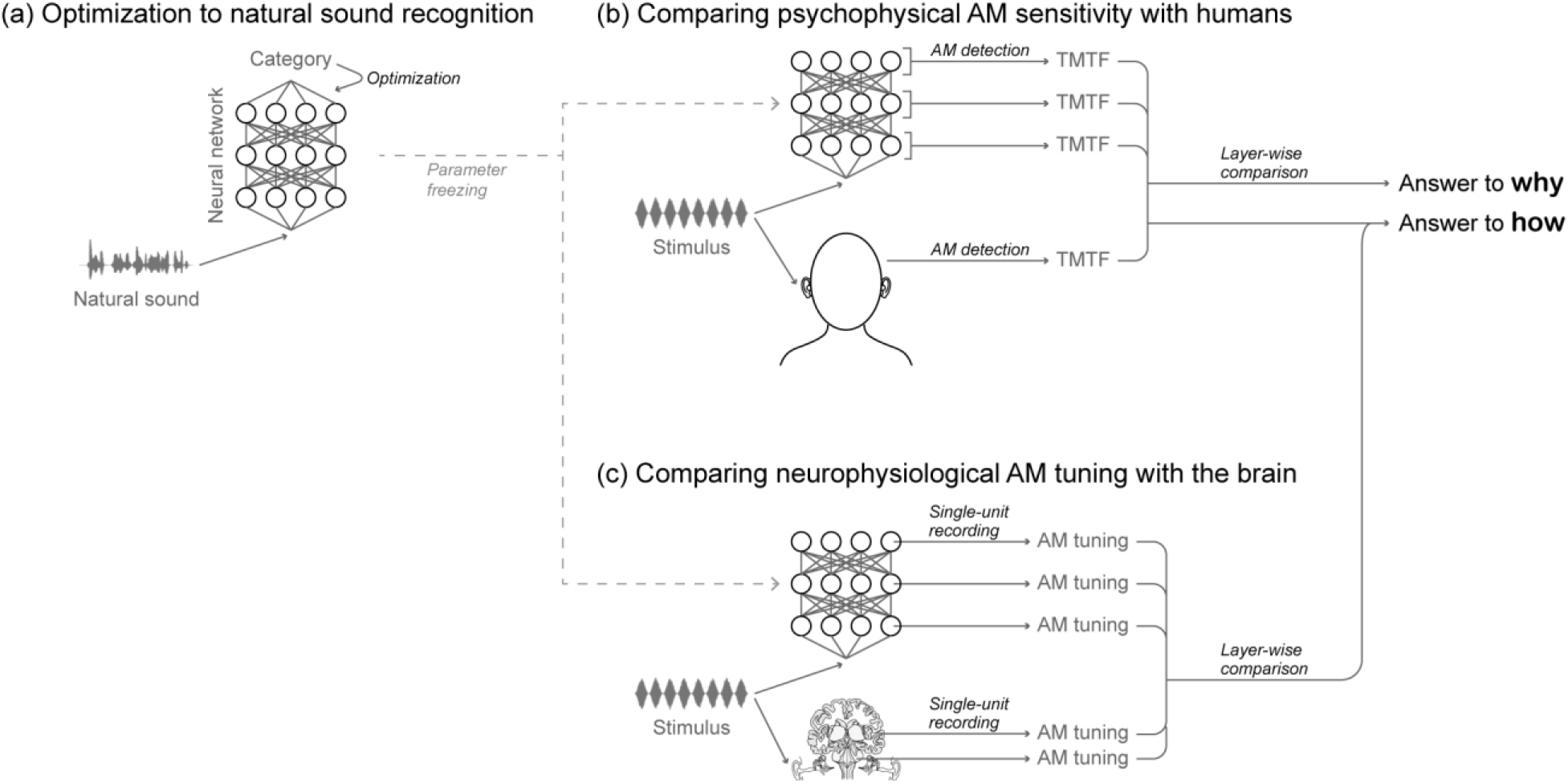
Schematic illustration of the framework of the present study, consisting of three stages (a)-(c). (a) Humans have evolved and developed the ability to precisely recognize natural sounds. We realized a computational simulation of this process by optimizing a model for natural sound recognition. Specifically, we used a deep NN that takes a sound waveform as input and estimates its category. (b) We froze the learned parameters and measured the AM sensitivity in the NN by using the same procedure as in human psychophysical experiments. A TMTF was computed for each layer. It was compared with previously reported human AM-sensitivity data in an attempt to answer why AM sensitivity has emerged in humans in its current form. (c) We measured neurophysiological AM tuning in the units in the NN by using the same procedure as in animal neurophysiological experiments. On the basis of the similarity of the AM tuning with the auditory brain regions and the results of the psychophysical experiments, we could infer possible neural mechanisms underlying behavioral AM sensitivity.

Finally, to investigate how AM sensitivity is realized, we performed neurophysiological experiments on the same optimized model and made a hierarchical correspondence with the ANS (Figure 3c). In parallel to the hierarchy of AM-tuned neurons identified by neurophysiology, our previous study found another hierarchy of AM-tuned units in the artificial NN (Koumura et al., 2019). By linking these two hierarchies, it established a method of layer-wise mapping of the AM representation between an NN and the ANS based on single-unit activities. Taking advantage of this neurophysiological layer mapping, in the current study, we roughly mapped the layer-wise AM sensitivity measured in the psychophysical simulation onto the hierarchical processing stages in the ANS. In this way, we could infer which auditory brain regions are most likely to be responsible for human AM sensitivity. Model optimization and all analyses regarding “why” and “how” were performed independently on two different sound recognition objectives. We believe that the finding of this study will contribute to our understanding of the complexity of the human auditory system by bridging the gap between the previous findings on psychophysical AM sensitivity in human behavior and neurophysiological AM tuning in the mammalian ANS.

Our very preliminary results have been published as a conference proceeding (Koumura, T., Terashima, H. & Furukawa, S. “Psychophysical” modulation transfer functions in a deep neural network trained for natural sound recognition. in Proceedings of the International Symposium on Auditory and Audiological Research 7, 157–164, 2020).

## Results

### Optimizing a neural network for natural sound recognition

Our NN consists of multiple layers, which in turn consist of multiple units (Figure 4). An input sound waveform was fed to the first layer, which performed temporal convolution and a static nonlinear operation. The outputs of the first layer were fed to the second layer, and this process continues to the topmost layer. There were no feedback or recurrent connections. Above the topmost layer was a classification layer that computed the categories of the input sound. The classification layer was not included in the psychophysical or neurophysiological analysis.

**Figure 4.**
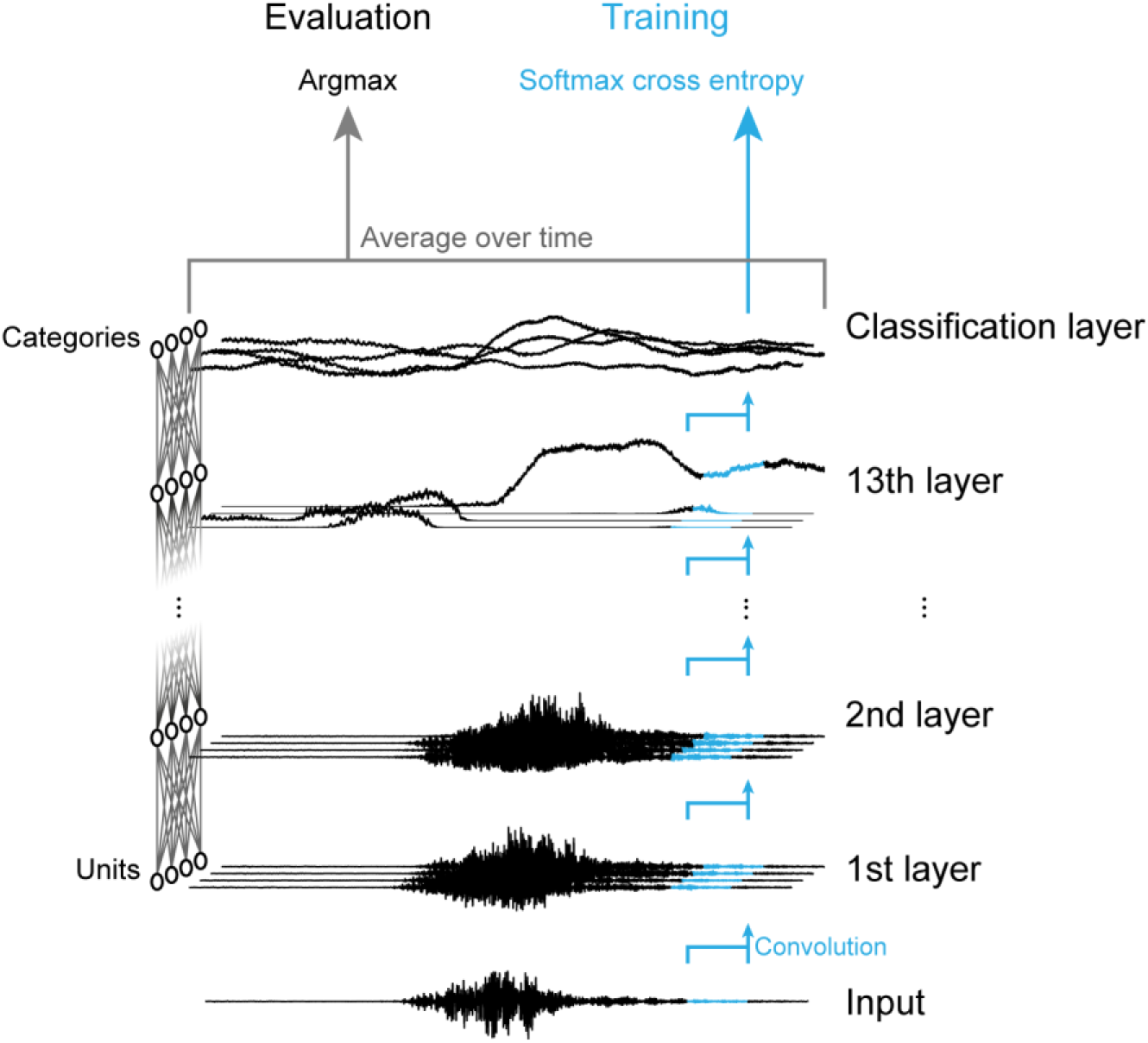
Schematic illustration of the NN architecture. Units in the first layer took a waveform as input and applied a non-linear temporal convolution to it. Subsequent layers took the activations in the layer below as input. Above the topmost convolution layer (13^th^ layer in the figure) was a classification layer. The number of units in the classification layer equals the number of sound categories. During training, softmax cross entropy was calculated for a single timeframe at a time (corresponding to the input sampling rate). During the evaluation, values in the classification layer were averaged over time and the category with the maximum average value was chosen as the estimated output category. The classification layer was not included in the psychophysical or neurophysiological analysis. This figure is a simplified illustration. The length of the convolutional filters and the number of units are not the same as those in the actual architectures used in this study.

The model was optimized to correctly classify natural sounds. We used two types of sounds, everyday sounds (Piczak, 2015) and speech sounds (Garofolo et al., 1993). The optimization objective was to correctly estimate the category of an everyday sound excerpt or the phoneme category at the center of a speech sound excerpt. We built and analyzed a model for each sound type. Because the results were generally consistent across different sound types, below we report the results for everyday sounds before those for speech sounds.

The recognition performance of an NN generally depends on its architecture (Bergstra and Bengio, 2012; Bergstra et al., 2013; Klein et al., 2017). In this study, we trained multiple NNs with different architectures and performed psychophysical and neurophysiological analyses on the NNs that achieved the highest recognition accuracy. To reduce possible biases by a specific architecture, unless otherwise stated, the reported recognition accuracy, TMTFs, and neurophysiological similarities are averages of the results of the four models with the highest recognition accuracies. This could be considered as a modeled version of reporting average quantities in multiple participants in human studies (Francl and McDermott, 2022). Four architectures with 13 layers were selected by performing an architecture search (see the methods for the detailed procedure). Their parameters are in Table 1.

**Table 1.** Architectural parameters of the models with the highest recognition accuracy. The layers all had the same number of units for simplicity. The size and dilation width of the convolutional filter were randomly sampled for each layer. The input time window of the filter was calculated as (dilation width) × (filter size – 1) + 1.

|  | Number of units per layer | Convolutional filter size (from lower to higher layers) | Dilation width (from lower to higher layers) |
| --- | --- | --- | --- |
| Architecture 1 | 256 | 5, 3, 6, 5, 2, 7, 8, 8, 7, 4, 4, 6, 5 | 231, 603, 18, 138, 14, 97, 105, 7, 137, 381, 193, 208, 266 |
| Architecture 2 | 512 | 3, 4, 4, 3, 6, 6, 4, 6, 8, 8, 7, 7, 3 | 449, 12, 161, 374, 175, 193, 120, 209, 161, 47, 151, 16, 465 |
| Architecture 3 | 128 | 7, 4, 4, 2, 7, 3, 7, 5, 2, 4, 3, 7, 2 | 42, 125, 341, 603, 96, 410, 44, 269, 747, 152, 528, 122, 823 |
| Architecture 4 | 512 | 3, 7, 2, 7, 7, 4, 5, 5, 3, 8, 8, 8, 2 | 676, 105, 581, 54, 16, 192, 214, 2, 173, 173, 184, 92, 887 |

After optimization, we evaluated the recognition performance for sounds not used in the model construction. The recognition accuracy was 0.477. This value is well above the chance level (0.02) but lower than those of state-of-the-art machine learning studies (Gong et al., 2021). Although tuning the hyperparameters or increasing the amount of training data may lead to an improvement in accuracy, we used the model as is in the subsequent analysis, because our goal was to understand the properties of the human hearing system, not to pursue accuracy improvements.

### Simulating psychophysical experiments as a way of measuring the models’ AM sensitivity

To investigate the relationship between sound recognition and AM sensitivity, we measured the TMTFs in each of the four best models by simulating psychophysical AM detection experiments. To fairly compare the model’s TMTF with those of humans, we replicated the procedures of human psychophysical experiments as precisely as possible. We simulated six psychophysical experiments from three independent human studies (Viemeister, 1979; Dau et al., 1997a; Lorenzi et al., 2001b, 2001a). In all of them, human subjects conducted a two- or three-interval forced-choice (2 or 3IFC) task. In each trial of the task, 2 or 3 stimuli were sequentially presented and only one among them was modulated. The task of a subject was to identify the modulated stimulus. At a given AM rate, an AM detection threshold was estimated with an adaptive method to find the AM depth that gave a 70.7% correct rate (Levitt, 1971).

The stimuli were sinusoidally amplitude-modulated broad- or narrow-band Gaussian noise. They differ in their stimulus parameters (Table 2). Because the most notable difference is in the carrier bandwidth (2 Hz, 3 Hz, 31 Hz, 314 Hz, or broadband), hereafter we will specify the conditions with their carrier bandwidths, except for two conditions with the broadband carrier. We call the broadband condition with the 0.5 s stimulus duration “broadband, short”, and the broadband condition with the 2 s duration “broadband, long”.

**Table 2.** Stimulus parameters in the AM detection experiments. Other parameters such as the fade duration vary among the studies but not all of them are shown here.

| Carrier bandwidth | Duration | AM starting phase | Amplitude equalization | Reference |
| --- | --- | --- | --- | --- |
| 2 Hz | 2 s | Random | After applying AM | Lorenzi 2001a |
| 3 Hz | 1 s | Constant | After applying AM | Dau 1997 |
| 31 Hz | 1 s | Constant | After applying AM | Dau 1997 |
| 314 Hz | 1 s | Constant | After applying AM | Dau 1997 |
| Broadband | 0.5 s | Constant | After applying AM | Viemeister 1979 |
| Broadband | 2 s | Random | Before applying AM | Lorenzi 2001b |

In the present study, to conduct an xIFC (x = 2 or 3) task, we presented a stimulus to the model and averaged the activities of each unit over the stimulus duration (Figure 5). Then, from the vector representing the units’ time-averaged activities in a single layer, we estimated the probability of the stimulus being modulated with logistic regression. The stimulus with the maximum probability was considered to be the model’s response to that xIFC trial. If the interval actually contained the modulated stimulus, the trial was considered correct. For simplicity, the threshold was estimated with a constant stimulus method. That is, the proportion of correct trials was computed independently for each AM depth. An asymmetric sigmoid function was fitted to the plot of the proportion of correct responses versus AM depth (Figure 5c). The threshold was defined as the AM depth at which the proportion correct was 70.7% on the fitted curve.

**Figure 5.**
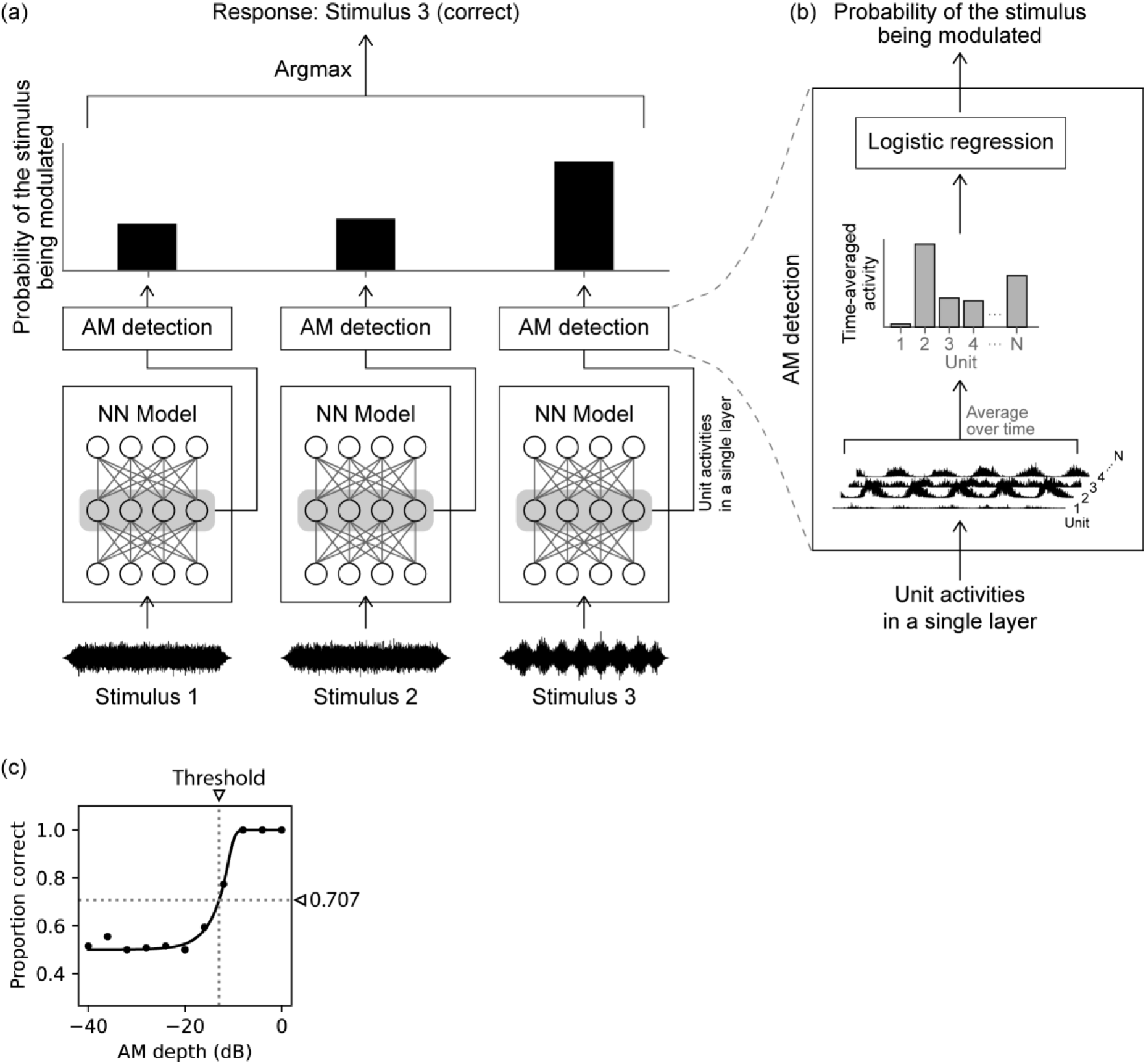
(a) Schematic illustration of the AM detection method in a 3IFC trial. Three stimuli were presented to the model and the probabilities of the stimuli being modulated were estimated for each layer from its unit activities. The probability was estimated independently for each stimulus. The interval with the maximum probability was taken to be the model’s response to the task. It was calculated for each layer. In this example, it is the third interval, which is correct because the third stimulus was modulated. (b) The boxes labeled “AM detection” in (a) are expanded for a detailed illustration of the probability estimation method. Logistic regression was applied to the time-averaged unit activities in a single layer. N denotes the number of units in the layer. (c) An example of a psychometric curve obtained from a single layer. The proportion of correct trials (dots) was fitted with an asymmetric sigmoid curve (solid line). The detection threshold was defined as the AM depth at a 0.707 correct proportion (dotted lines).

### Emergence of human-like TMTFs in the model

The forms of the model’s TMTFs (detection thresholds as a function of AM rate) varied depending on the stimulus condition and model layer (Figure 6, orange lines). The forms of the human TMTFs (black dotted lines) also depend largely on the stimulus condition. In all conditions, the model and human TMTFs tended to overlap in the middle to higher layers.

**Figure 6.**
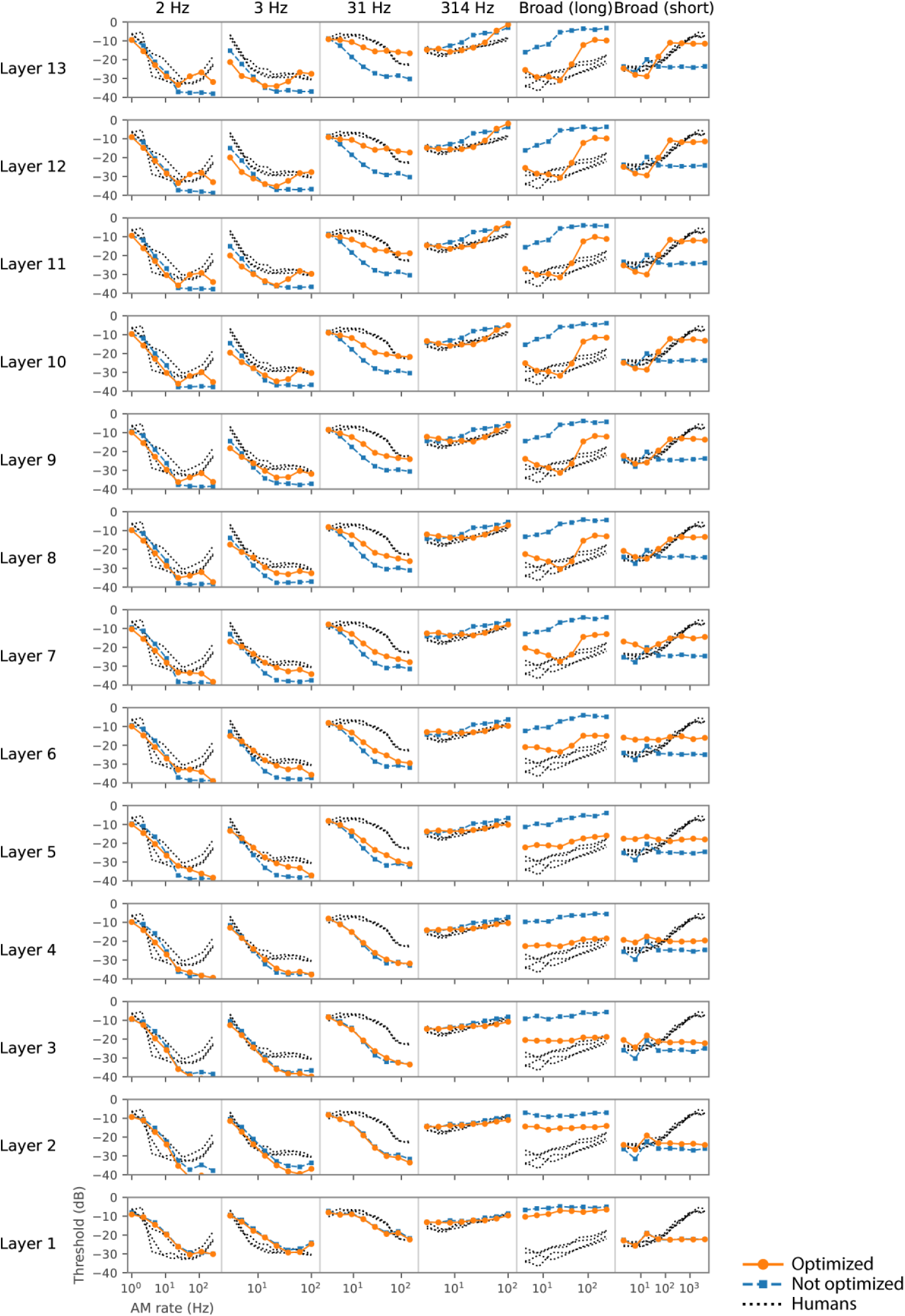
TMTFs in the model optimized to everyday sounds (orange circles), those in the non-optimized model (blue squares), and those in humans (black dotted lines). The columns correspond to different experimental conditions, while the rows correspond to the different layers. The TMTFs in the higher layers of the optimized model appear to be more similar to those of humans than those of the lower layers or the non-optimized model.

Quantitative analyses supported the above observations. The similarity of the model’s TMTFs with humans’ was evaluated in terms of relative patterns (reflecting mainly similarity in TMTF shape) and absolute values (reflecting similarity in both shape and sensitivity in dB) (Figure 7). An index of similarity of relative patterns, the correlation coefficient, was calculated from pairs of model and human TMTFs. Hereafter, we call it the *pattern-similarity index* (Figure 7, upper panel). As an index of absolute measure of dissimilarity, we calculated the root mean square (RMS) deviation and called it the *discrepancy index* (Figure 7, lower panel). To take all stimulus conditions into account, TMTFs in all stimulus conditions were pooled when calculating the indices. The two measures consistently indicated that layers around the 10th layer exhibited TMTFs most similar to humans’ (highest pattern similarity and lowest discrepancy). This result indicates the emergence of human-like AM sensitivity in the model optimized for natural sound recognition.

**Figure 7.**
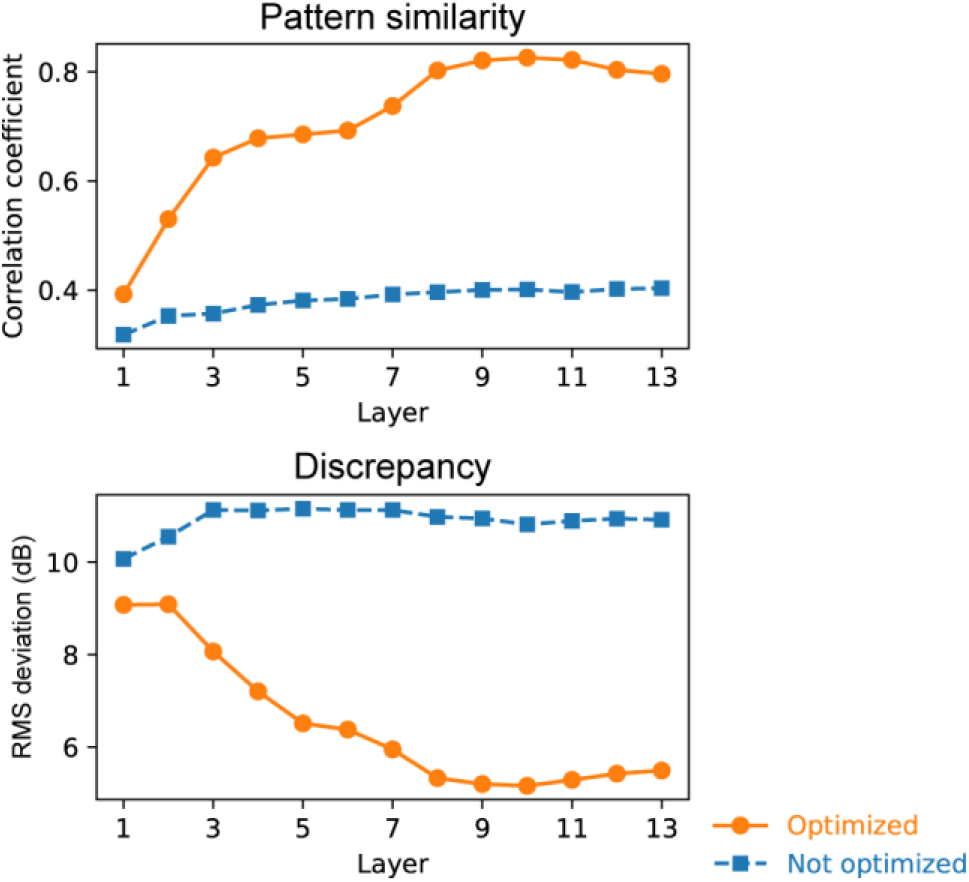
Quantitative comparison of the TMTFs in the model and humans. Pattern-similarity index (top panel) and discrepancy index (bottom panel) in the models optimized to everyday sounds (orange circles) and the non-optimized models (blue squares) are shown. The relatively higher layers of the optimized models show large pattern similarity and small discrepancy. The lower layers and the non-optimized models show low similarity.

Human-like TMTFs did not emerge in the non-optimized model with random initial parameters. The sound recognition accuracy in the non-optimized model was 0.013, which was as low as the chance level, 0.02. Generally, the TMTFs in the non-optimized model were relatively invariant across the layers and showed marked discrepancies from those of humans and the optimized models. These discrepancies were particularly apparent in the higher layers (Figure 6). These observations are supported by the quantitative analyses, showing a low pattern-similarity index and high discrepancy index throughout the layers (Figure 7). These results suggest that optimization to sound recognition is an essential factor for the emergence of human-like TMTFs, and that the NN architecture only could not explain human AM sensitivity.

### Models with better recognition performance were more human-like

An additional analysis revealed a close link between the model’s sound recognition performance and the TMTF similarities. During the NN architecture search, we trained 20 models with 13 layers with different architectures (remember that the above analyses targeted the best four models of the 20). The 20 models exhibited recognition accuracies ranging from 0.043 to 0.492 and produced TMTFs with a varying degree of similarity to humans (Figure 8a, Figure 8b). We examined the relationship between the recognition accuracy and TMTF similarity indices (i.e., pattern-similarity and discrepancy indices).

**Figure 8.**
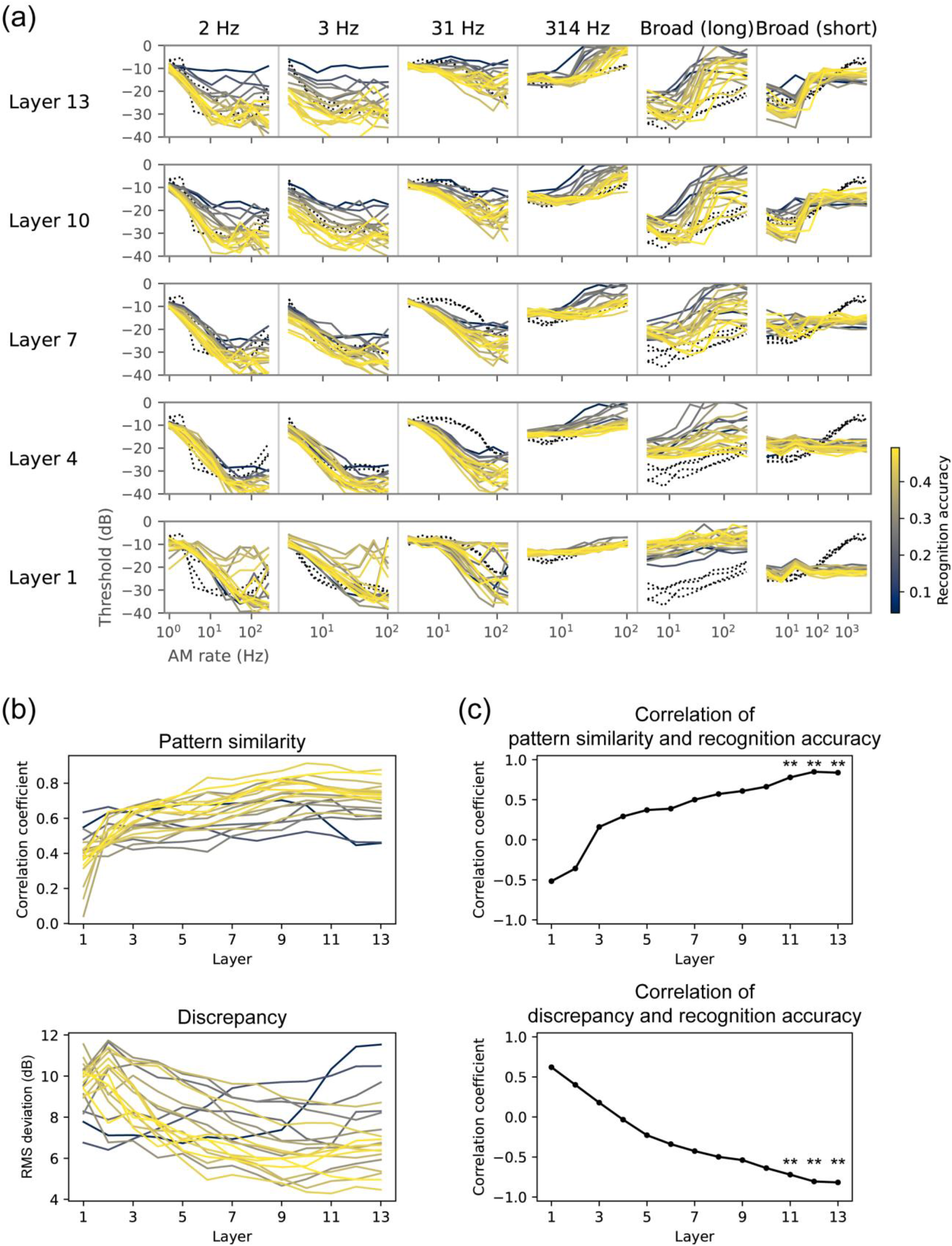
AM sensitivity in different architectures and its relationship to recognition performance. (a) TMTFs in the models with different architectures. Each colored line shows results for a single model with a specific choice of NN architecture. The color indicates the recognition accuracy of the corresponding architecture. Black dotted lines show human TMTFs. (b) Pattern similarity and discrepancy indices. (c) Correlation coefficients between the (dis)similarity indices and the recognition accuracy. Statistically significant positive and negative correlations were found in the highest layers. ** p < 0.01 with a Bonferroni correction for the number of layers.

The TMTF similarity indices correlated with the recognition accuracy in the higher layers (Figure 8c). The positive and negative correlations, respectively for pattern-similarity and discrepancy indices, mean that the models that performed sound recognition better exhibited AM sensitivity more similar to that of humans. This result further supports the idea that there is a strong relationship between optimization for natural sound recognition and emergent AM sensitivity.

### Training signals must have natural AM patterns for the emergence of human-like AM sensitivity

What components of the optimization for natural sound recognition are essential for acquiring human-like AM sensitivity? We hypothesized that natural AM patterns in the sound are the critical feature. To test this hypothesis, we conducted control experiments in the models optimized for the manipulated sound signals. Only the training signals were manipulated. No modifications were made to the stimuli for measuring AM sensitivity.

The manipulation involved dividing a sound signal into an amplitude envelope (Env) and temporal fine structure (TFS) and disrupting either of them while preserving the other, for the entire signal or each sub-band. This is a common strategy for manipulating the AM structure of sound in auditory science (Smith et al., 2002; Lorenzi et al., 2006). Specifically, we tested the following four types of signals (see methods for details):

1. *Single-band Env signals*, which preserved Env while TFS was replaced with that of broadband noise in the entire signal.
2. *Multi-band Env signals*, in which the original signal was divided into multiple frequency bands; Env for each band was preserved while TFS was replaced with a random narrowband noise corresponding to that band; then, the multiband signals were added together.
3. *Single-band TFS signals*, which, for the entire signal, preserved TFS while Env was flattened.
4. *Multi-band TFS signals*, in which the original signal was divided into multiple frequency bands; TFS for each band was preserved while Env was flattened; then, the multiband signals were added together.

We optimized the NN models to recognize the manipulated sounds. Hereafter, we call the optimized models for the above signals the *single-band Env model, multi-band Env model, single-band TFS model*, and *multi-band TFS model*. We refer to the model trained for intact sounds (i.e., the one described in the earlier sections) as the original model.

Differences in the TMTFs across the models were more apparent for higher layers. TMTFs of the single-band ENV model appeared to be closest in general shape to the human TMTFs (Figure 9a, blue lines). The pattern-similarity index for the single-band Env model was at a comparable level to the original model throughout its layers (Figure 9b top). However, the discrepancy index of the single-band Env model deviated from the original one in the layers above the 8^th^, exhibiting higher values (Figure 9b middle). These results indicate that the model had TMTFs whose patterns were similar to humans, while its sensitivity to AM was higher than that of humans (i.e., lower thresholds; the blue lines were generally lower than the black dotted lines, Figure 9a). This difference in AM sensitivity was quantified as the *net difference*, the average signed difference between the model TMTFs and the human TMTFs (Figure 9b bottom). The net difference was largely negative in the single-band Env model, indicating that its thresholds were on average lower than humans’. These results suggest that optimizing to sounds that only retain natural single-band AM patterns made the model more sensitive than humans to AM. This is probably because the model was biased toward exploiting the AM that was the only available feature for recognition.

**Figure 9.**
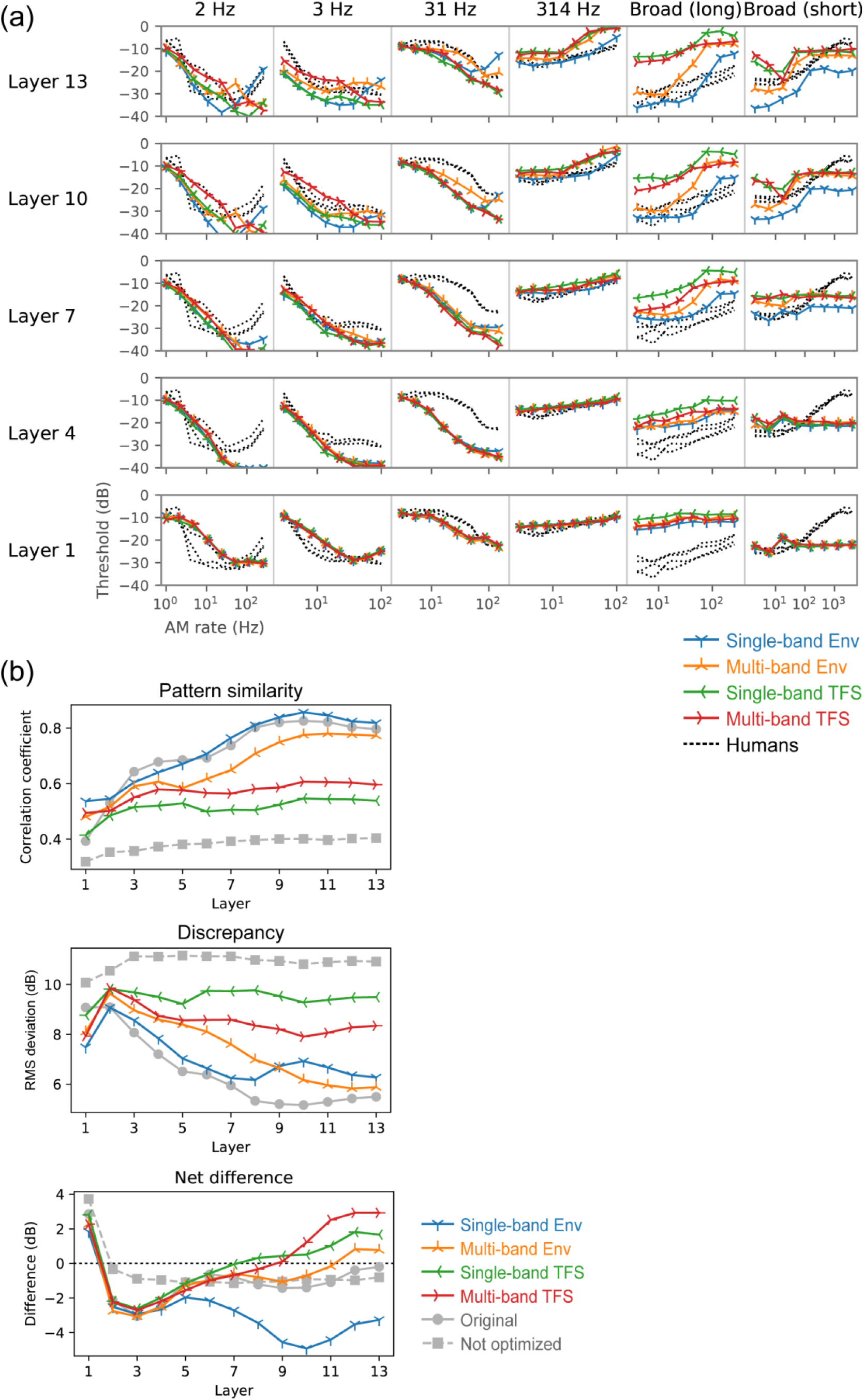
(a) TMTFs of the models optimized to degraded sounds. (b) Their pattern-similarity index (top panel), discrepancy (middle panel), and net difference from humans (bottom panel). The indices of the original optimized and non-optimized models are shown as grey lines. Overall, in the higher layers, the TMTFs of the Env models were more similar to those of humans than the TMTFs of the TFS models were. Single-band Env models exhibited high pattern similarity but also showed a high discrepancy, indicating that the patterns of the TMTFs, but not their absolute values, were similar to those of humans. Their thresholds appeared to be lower than those of humans, as shown by the negative net difference.

The TMTFs of the multi-band Env model were somewhat similar to those of humans. Both pattern-similarity and discrepancy indices gradually approached the original model with increasing layer number, reaching a comparable level at the 12th and 13th layers.

In contrast, the TMTFs of the single-band and multi-band TFS models were consistently different from those of humans. This result, together with the results of the two Env models, suggests that natural AM patterns are essential for the emergence of human-like AM sensitivity.

It is important to note that all models achieved sufficiently high accuracy in the sound recognition task, well above chance level, and that the accuracies of the single-band Env and TFS models were comparable (Figure 10). This indicates that all the signals contained a sufficient amount of information for sound recognition and that all the models were capable of using the information. Thus, the non-human-like TMTFs of the TFS models could not be attributed to failures in their optimization.

**Figure 10.**
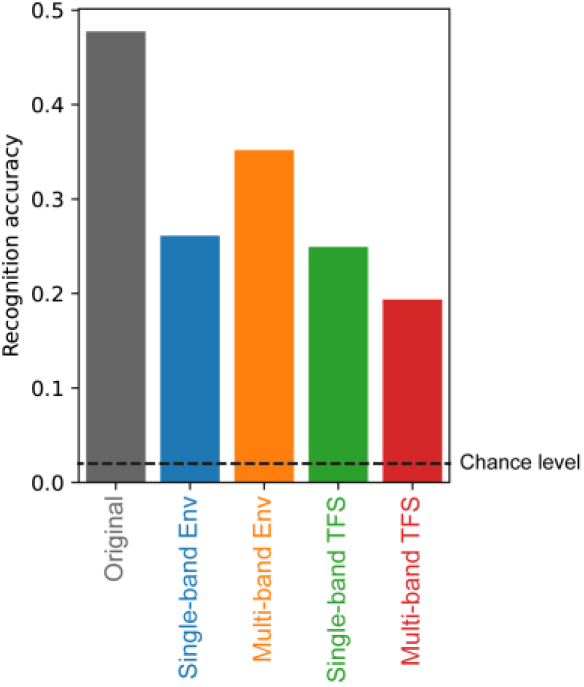
Recognition accuracy of models optimized to degraded sounds. The result of a model optimized to the original sounds is also shown on the left. Generally, the model’s recognition accuracy dropped when it was optimized to degraded sounds, but the drop was not catastrophic.

### Comparison with temporal-template-based AM detection strategy

In this study, we assumed AM detection is based on time-averaged activities in the model. On the other hand, previous computational studies used a method based on the temporal-template for simulating the psychophysical AM detecting process (Dau et al., 1997a). There might be a possibility that different detection strategies yield different forms of TMTF. In their study, template-based AM detection was performed on the outputs of the MFB. A template for a given AM rate was generated by averaging the MFB outputs over multiple independent carrier instances. In an xIFC trial, a correlation coefficient was calculated between the template and the MFB output for a test signal. The stimulus interval with the highest correlation was chosen as the response of the model. Here, to test whether the same detection method works well for our NN model, we applied it to the unit activities in a single NN layer (Figure 11a). A template was defined as the average unit activities in response to fully modulated stimuli minus the average activities in response to non-modulated stimuli. The average was taken over multiple carrier instances. Then, in each trial of the xIFC task, the model’s response was defined as the stimulus interval with the largest correlation between the model’s activities and the template. In accordance with the previous study (Dau et al., 1997a), the correlation was non-normalized, i.e., the dot product.

**Figure 11.**
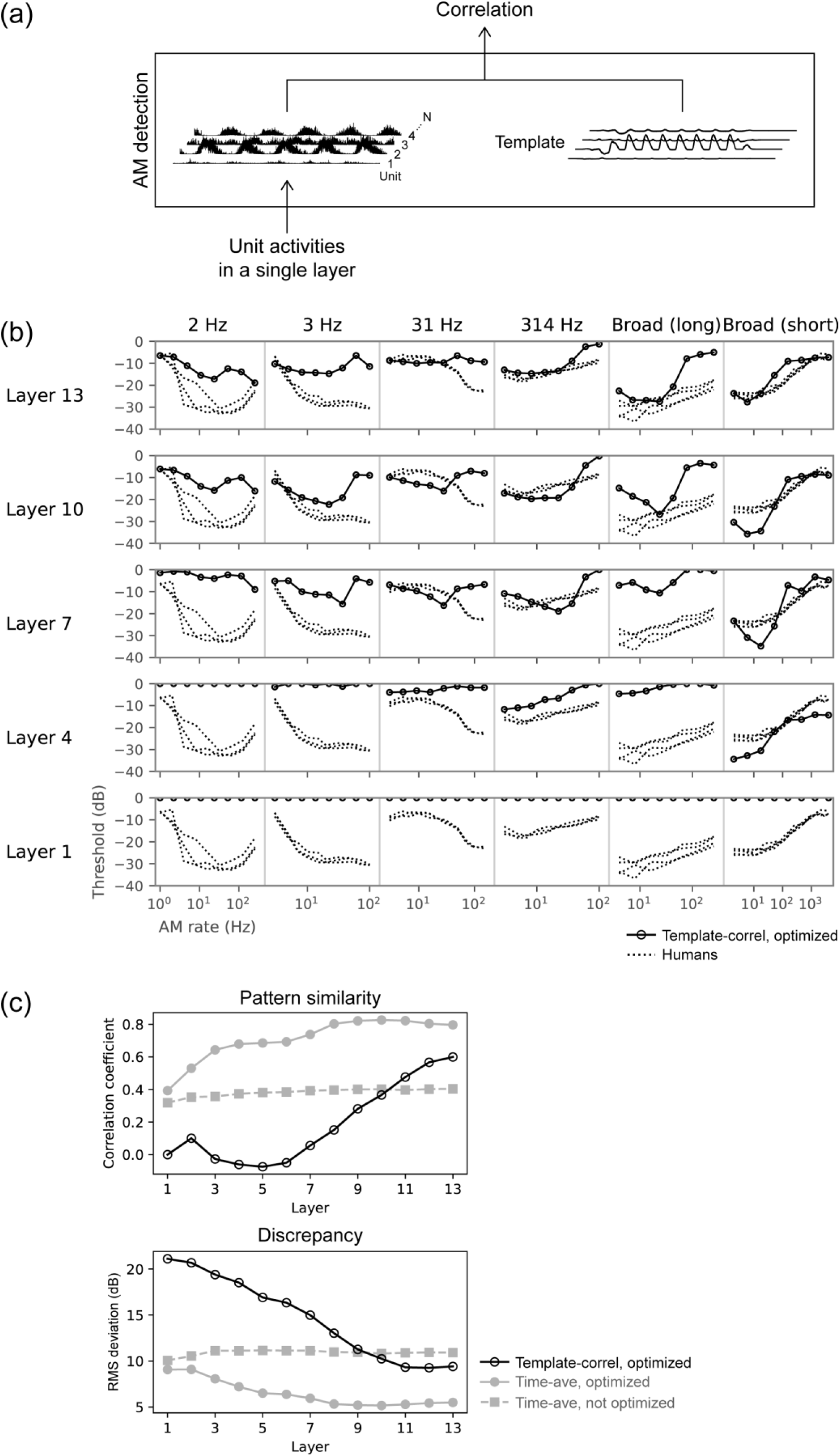
(a) Schematic illustration of AM detection process based on correlation with a template. For purpose of explanation, this illustration replaces Figure 5b, where the correlation in this figure corresponds to the output probability in Figure 5b. (b) TMTFs obtained from AM detection based on template correlation (open circles). TMTFs of humans are shown as dotted lines. (c) Pattern-similarity index and discrepancy index from the template-based detector (black open circles). The similarity indices for the time-average-based detector (Figure 7) are shown as filled symbols. AM detection based on template correlation did not result in human-like TMTFs.

The resulting TMTFs differed from human TMTFs (Figure 11b). The similarity to human TMTFs was the highest at the topmost layers (Figure 11c, open circles), but it was still lower than the similarity of the TMTFs of the time-average-based detector (Figure 11c, grey circles; same data as in Figure 7). This result indicates that human-like AM sensitivity was not observed in our model when it used temporal correlation with templates for AM detection. Template-based detection might work well for MFB outputs, but not for NN activities. Our results alone could not elucidate the reason for this difference. Perhaps different AM detection strategies should be applied to different sound representations (in an NN or in an MFB output).

### Neurophysiology of the model suggested involvement of the auditory midbrain and higher regions

In an earlier section, we indicated that human-like AM sensitivity was observed when the stimulus representation in layers around the 10^th^ layer was used for AM detection (Figure 7). Does this finding provide a significant insight into neural processes in the ANS? More specifically, to which brain regions do these layers correspond?

We addressed this question by mapping the NN layers onto the auditory brain regions based on the similarity of their neurophysiological AM tuning, as in our previous study (Koumura et al., 2019). Neurophysiological AM tuning in the model was measured by simulating neurophysiological experiments. The neuronal tuning properties in the ANS were taken from the neurophysiological literature (Müller-Preuss, 1986; Langner and Schreiner, 1988; Schreiner and Urbas, 1988; Batra et al., 1989; Preuß and Müller-Preuss, 1990; Frisina et al., 1990; Joris and Yin, 1998, 1992; Rhode and Greenberg, 1994; Zhao and Liang, 1995; Bieser and Müller-Preuss, 1996; Condon et al., 1996; Schulze and Langner, 1997; Eggermont, 1998; Huffman et al., 1998; Joris and Smith, 1998; Kuwada and Batra, 1999; Krishna and Semple, 2000; Lu and Wang, 2000; Lu et al., 2001; Liang et al., 2002; Batra, 2006; Zhang and Kelly, 2006; Bartlett and Wang, 2007; Scott et al., 2011; Yin et al., 2011). The target brain regions from peripheral to central were auditory nerves (AN), the cochlear nucleus (CN), superior olivary complex (SOC), nuclei of the lateral lemniscus (NLL), inferior colliculus (IC), medial geniculate body (MGB), and auditory cortex (AC).

The neurophysiological similarity of the NN layers and brain regions shows that lower and higher layers were relatively similar to the peripheral and central brain regions, respectively (Figure 12). Thus, the results of our previous study were replicated with the newly constructed model. Layers around the 10th layer roughly corresponded to the IC, MGB, and AC. This result indicates that the NN layers that exhibited human-like AM sensitivity had similar neural representations to those of the IC and higher brain regions.

**Figure 12.**
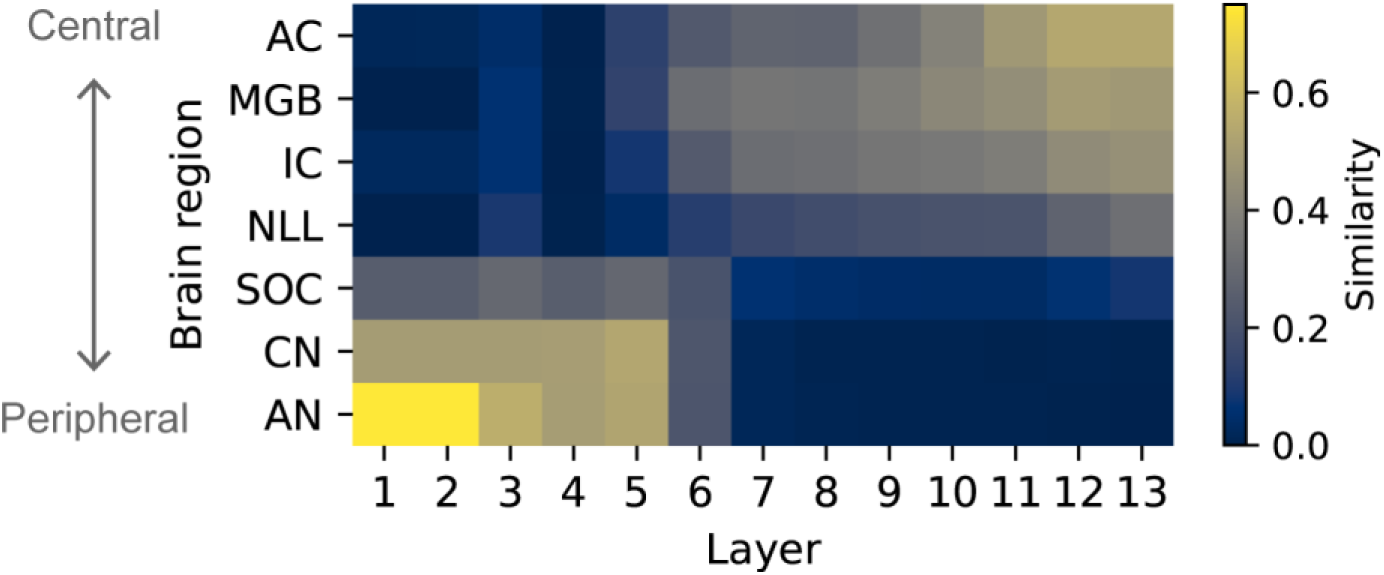
Similarity of the neurophysiological tuning between brain regions and NN layers. Layers that showed similar TMTFs with those of humans roughly correspond to higher regions like the IC, MGB, and AC. Abbreviations: AN, auditory nerve; CN, cochlear nucleus; SOC, superior olivary complex; NNL, nuclei of the lateral lemniscus; IC, inferior colliculus; MGB, medial geniculate body, and AC, auditory cortex.

### Emergent TMTFs through optimization to speech sounds

We conducted the same analysis on the models optimized for phoneme classification of speech sounds (Garofolo et al., 1993). The results were generally consistent with those of the models optimized for every day sounds reported above. This indicates that human-like TMTFs robustly emerged in the models that were independently optimized to two different types of sound.

The phoneme classification accuracy was 0.747, which is high above the chance level, 0.026. The TMTFs of the layers around the 8th and 9th layers in the optimized model were similar to those of humans (Figure 13a and b, orange lines), whereas neither the TMTFs of the non-optimized model nor those calculated with the template correlation were similar. According to the neurophysiological analysis, the brain region most similar to the 8^th^ and 9^th^ layers was the IC (Figure 13c). The MGB and AC also exhibited high neurophysiological similarity.

**Figure 13.**
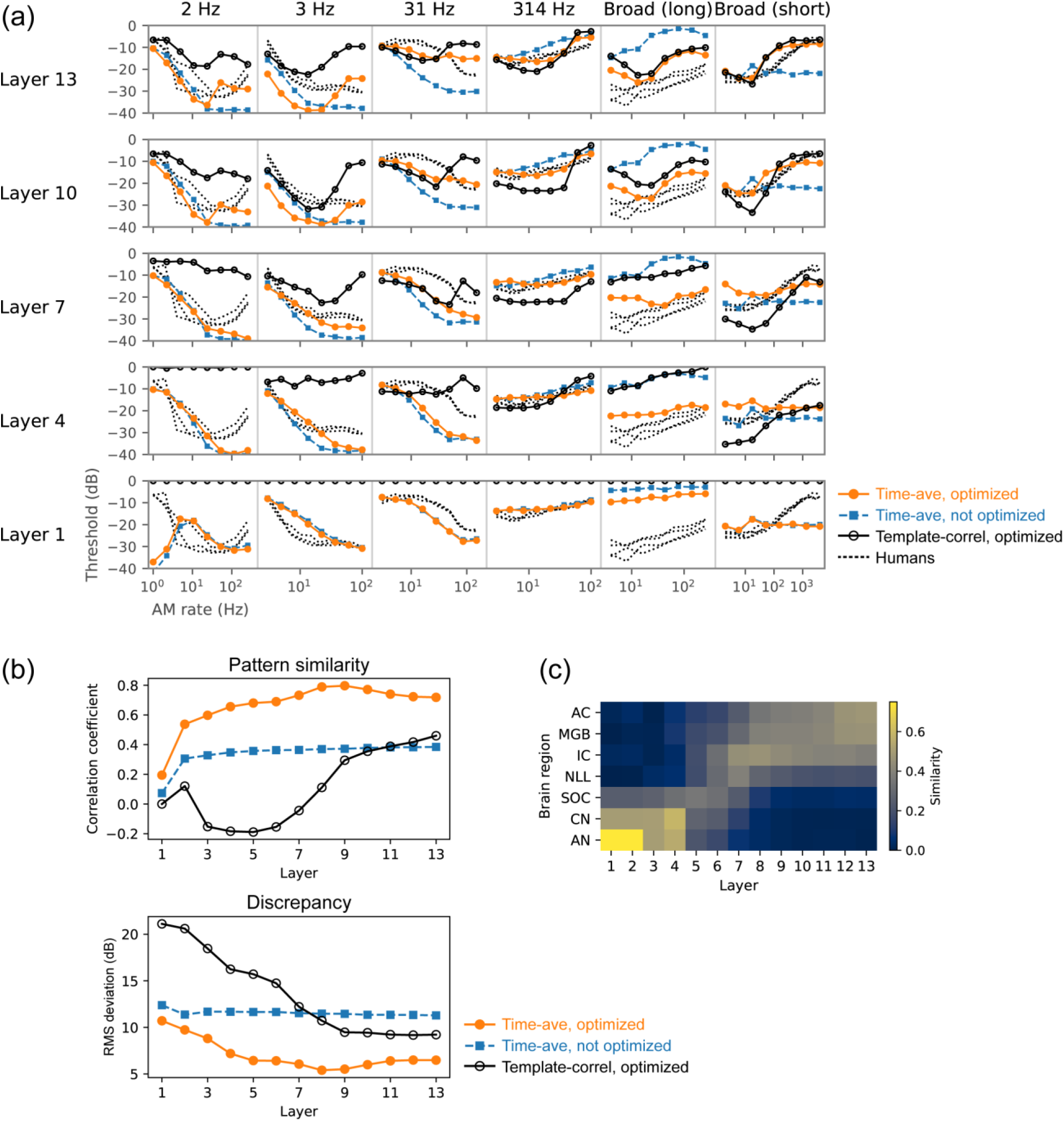
Results of the model optimized to speech sounds. (a) TMTFs of the optimized model with the AM detection process based on time-averaged unit activities (orange circles), those of the non-optimized model (blue squares), those from AM detection based on temporal correlation with templates (black open circles), and those of humans (black dotted lines). (b) Pattern-similarity index and discrepancy index between the model TMTFs and human TMTFs. (c) Similarity of neurophysiological tuning between NN layers and auditory brain regions.

On the other hand, we did not observe high correlations between similarity to human TMTFs and recognition accuracy. The pattern-similarity indices and the discrepancy indices did not exhibit appreciable variations across the 20 models with different architectures (Figure 14a), and there was no significant correlation with recognition accuracy (Figure 14b). This lack of correlation may be explained by the dynamic range of the recognition accuracy in the models for speech sounds: Their recognition accuracies ranged from 0.499 to 0.771, whereas, for the models optimized to everyday sounds, the range was from 0.043 to 0.492. All models trained on speech sounds showed recognition accuracies well above chance level, whereas some models trained on everyday sounds exhibited very low recognition accuracy almost as low as chance level. Probably, the correlations between the recognition accuracy and the similarity to human TMTFs are non-linear. Models that failed to perform sound recognition exhibited non-human-like AM sensitivity, but models with good sound recognition performance exhibited more-or-less human-like AM sensitivity regardless of the small variation in their performance. Probably, once the performance of sound recognition surpasses a certain level, the similarity of AM sensitivity to humans does not change largely upon any further increase in recognition accuracy.

**Figure 14.**
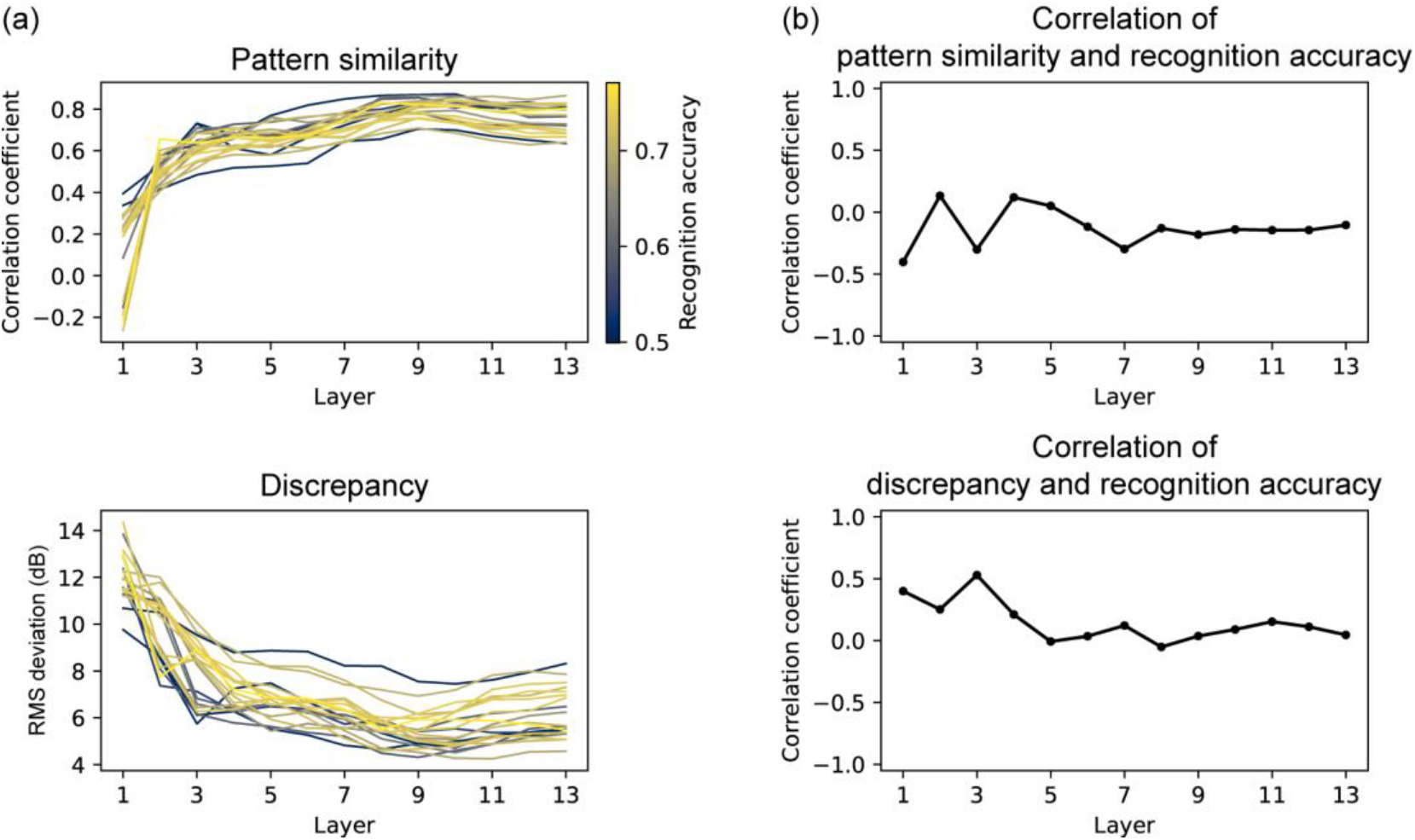
(a) Pattern similarity and discrepancy indexes of the TMTFs in the models with different architectures optimized to speech sounds. The pattern of the similarity indexes appeared similar across different NN architectures. (b) Correlation between similarity indexes and recognition accuracy. Significant correlation was not observed. These results are probably because of the small dynamic range of recognition accuracy.

When the Env or TFS was disrupted in the speech sounds, the recognition accuracies of the optimized models were well above the chance level, except for the multi-band TFS model (Figure 15a). The multi-band Env model had the highest recognition accuracy, followed in order by the single-band TFS model, single-band Env model, and multi-band TFS model. This order is consistent with human performance as shown in a previous study (Smith et al., 2002) and thus supports the conclusion that our models behave similarly to humans when recognizing those degraded speech sounds. The multi-band TFS model trained on everyday sounds showed relatively low recognition accuracy, although it was above the chance level (Figure 10).

**Figure 15.**
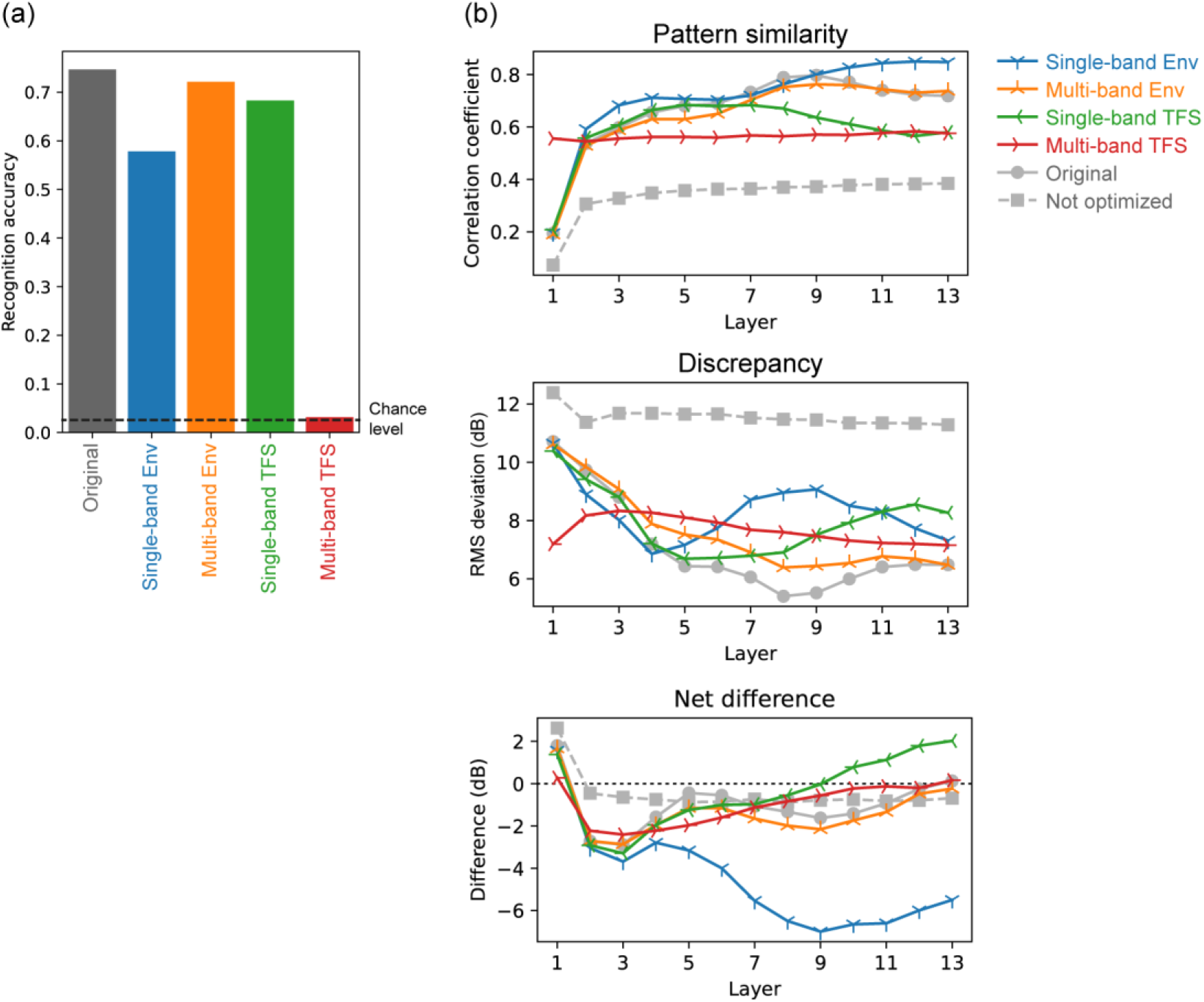
(a) Recognition accuracy of the models optimized to degraded speech sounds. The result for the model optimized to original speech sounds is also shown on the left. Recognition accuracy dropped when the models were optimized to degraded sounds, but the drop was not catastrophic except in the multi-band TFS model. (b) Pattern-similarity indices (top panel), discrepancy indices (middle panel), and net difference (bottom panel) between model TMTFs and human TMTFs. The results are consistent with those of the models optimized to everyday sounds.

The TMTFs of the models optimized to degraded speech sounds were qualitatively consistent with those of the models optimized to degraded everyday sounds (Figure 15b). The TMTFs of the single-band Env model exhibited high pattern similarity to those of humans, and their net difference shows their thresholds were lower than those of humans. The TMTFs of the multi-band Env model were similar to those of humans in terms of both their pattern and absolute value. The TMTFs in the TFS models were not similar to humans’.

## Discussion

### AM sensitivity emerging through sound recognition

The present study demonstrated that a model optimized to natural sound recognition exhibited human-like AM sensitivity. It should be emphasized that, in building the model, we did not make any attempt to design model architectures or adjust parameters for achieving closer similarity to humans, nor did we use any knowledge about human AM sensitivity. The results therefore suggest that the nature of AM sensitivity in humans might also be a consequence of optimizing to natural sound recognition in the course of human evolution and/or development in the natural environment. This notion is supported by the finding that natural AM patterns were essential for the emergence of human-like AM sensitivity.

By simulating previous human experiments as precisely as possible, we could quantitatively compare the TMTFs of our model and of humans. We were able to explain TMTFs in six experiments conducted in three independent human studies with a single unified modeling framework. Our results indicated that detailed patterns of AM sensitivity (i.e., TMTFs) have significant implications in natural sound processing. Our findings strengthen the existing knowledge that general AM sensitivity is closely linked to sound recognition ability (Cazals et al., 1994; Fu, 2002; Luo et al., 2008; Won et al., 2011; De Ruiter et al., 2015; Bernstein et al., 2016).

We built two types of models using either everyday sounds or speech sounds and analyzed each one independently. The results on the two datasets were qualitatively similar, although some relationships were stronger for everyday sounds. They suggested that human-like AM sensitivity is related to both sound types. This conclusion is consistent with the previous studies on cochlear frequency tuning and neural AM tuning, where qualitatively similar tunings were obtained from optimization to everyday sounds and speech sounds (Smith and Lewicki, 2006; Koumura et al., 2019). There might be a common representation of these sound types in the auditory system perhaps because the human auditory system has taken advantage of already evolved mechanisms to represent everyday sounds and built speech recognition functions on top of it.

### Relation to modulation filterbank theory

A number of psychoacoustic phenomena, including human TMTFs, have been explained by computational models which incorporate the MFB (Jepsen et al., 2008). The MFB is a conceptual realization of midbrain neurons that are tuned to the modulation rate. It was formalized to explain various psychophysical phenomena, e.g., masking in the modulation domain (Dau et al., 1997a, 1997b). It should be noted that our NN model, in contrast, does not include any explicit implementation of auditory mechanisms (e.g., a cochlear filterbank or an MFB), nor does it attempt to reproduce any psychophysical phenomena (e.g., modulation masking). This is because the purpose of our study is not delving into the signal processing mechanisms of the AM sensitivity, but to investigate emergent TMTFs in a sound recognizing model and the effect of optimization and sound features to be optimized on the emergent AM sensitivity. Thus, it is difficult to make a fair comparison between models based on MFB theory and those in the present study. Nevertheless, we consider it worth discussing the present findings in relation to MFB theory.

Compared with the TMTFs in previous computational studies involving the MFB model (Jepsen et al., 2008), the TMTFs of our model were less similar to human TMTFs. The TMTFs in Jepsen et al. 2008 showed a pattern-similarity index of 0.95 and discrepancy index of 2.3 dB (derived from FIG. 8 in Jepsen et al. 2008; caveat: they did not test a 2-Hz bandwidth or long-broadband conditions). This, however, is not surprising and does not indicate the inferiority of our model study. That is, the previous study designed the model explicitly for reproducing human psychophysical properties including AM sensitivity, whereas our model does not explicitly try to reproduce any psychophysical or neurophysiological properties in the auditory system.

The major components of the MFB model include frequency decomposition by a cochlear filterbank, subsequent amplitude envelope extraction, and an MFB running on the envelope of each frequency band. Our NN model does not include any of these implementations explicitly. Nevertheless, we can infer that the signal processing in our model is in line with MFB theory for the following reasons. First, our previous study has shown that middle layers in NNs exhibit units with tuning to various AM rates, which probably work as bandpass filters with various frequency responses (Koumura et al., 2019). Second, convolution itself can work as a bandpass filter and a stack of convolutions with static nonlinearity can work as an envelope extractor. Such a stack can extract the signal envelopes and then apply a bandpass filterbank to them. Therefore, our model may have implicitly acquired a function equivalent to an MFB in the course of optimization to natural sound recognition.

Whether or not to apply additive internal noise is another critical difference between the models. In the MFB models (e.g., (Dau et al., 1997a)), additive noise was applied to the model’s internal representation from which AM detection was simulated. In contrast, our model is deterministic, meaning that the unit responses are the same for the same stimulus, except for the non-deterministic behaviors of atomic operations in a GPU. In the AM detection experiments on our model, the only source of stochasticity was the stimulus, due to the noise carrier and the AM starting phase in the conditions with a random starting phase, both of which were sampled independently stimulus by stimulus (Table 2). Psychophysically relevant internal noise (e.g., as proposed by (Ewert and Dau, 2004)) could increase the similarity of the model’s TMTFs to human ones.

### Neural mechanisms of behavioral AM sensitivity

#### Hierarchical brain regions and AM detection

Layer-wise measurement of TMTFs revealed different AM sensitivities along the layer cascade. In the optimized model, TMTFs in the middle to higher layers were most similar to humans’. Here, we discuss the implications of this result in terms of signal processing and anatomical brain regions.

Generally, an optimized NN behaves as an effective signal processor and feature extractor. While processing an input signal, each of its cascading layers computes its representation by integrating and non-linearly transforming the one below. Its first layer computes relatively simple and temporally and/or spatially local features directly from the input. As the processing stage progresses, the extracted features gradually become more complex and global (Mahendran and Vedaldi, 2015; Yosinski et al., 2015) and a representation that is effective for input recognition gradually emerges (Carter et al., 2019; Bau et al., 2020). Therefore, our results suggest that the AM detection ability of humans might be based on relatively higher-order features of the stimulus.

Interestingly, this tendency throughout the processing hierarchy can also be seen in the ANS. Peripheral regions in the auditory pathway are generally sensitive to fast temporal changes in a sound signal and relatively linear features, whereas central regions are sensitive to slower changes and more non-linear features (Joris et al., 2004; Sharpee et al., 2011). In the present study, as well as calculating the psychophysical AM sensitivity in the NN layers, we also calculated the neurophysiological similarity between the NN layers and different brain regions. We found that layers with human-like psychophysical TMTFs showed neurophysiological AM tuning similar to that in the IC, MGB, and AC (Figure 12, Figure 13c). This result suggests that human-like TMTFs could be observed when conducting an AM detection task from the stimulus representation in these brain regions. A human brain might also use stimulus representations in the IC, MGB, and AC when conducting AM detection tasks, but our results alone cannot distinguish whether such a computation is running within those brain regions or somewhere outside, possibly regions associated with higher-order cognitive functions. Another unanswered question is which (possibly all?) of these regions is actually the source of the neural representation used by the AM detector (if it exists) implemented in the human brain. Nevertheless, the present results, at least, suggest that the stimulus representation necessary for AM detection emerges as early as in the IC and is kept until the signal reaches the AC.

#### Neural AM representation relevant to AM detection

We obtained human-like TMTFs by assuming an AM detection process based on time-averaged unit activities. Time-averaged activities in a unit can be interpreted as the average firing rate of a neuron (Koumura et al., 2019). Taken together with the above discussion, it can be suggested that the behavioral AM sensitivity of humans might be based on the average firing rate in the IC, MGB, and AC. This is consistent with the previous neurophysiological findings that relatively central auditory brain regions, such as the IC, MGB, and AC, perform rate coding of AM (Joris et al., 2004), which means that their average firing rate encodes the stimulus AM rate.

The other AM coding strategy in the ANS is temporal coding, in which temporal patterns of neural activities encode the AM rate (Joris et al., 2004). We also tested temporal-coding based AM detection by performing the temporal-template-correlation method, but findings were elusive. Our results alone could not distinguish whether or not the brain relies on temporal coding when performing AM detection. There is a possibility that our template-based detection strategy is too simple to simulate a temporal-coding-based AM detection process in the human brain. Using a more sophisticated detection process, such as learnable convolutional mapping (Bashivan et al., 2019), might result in more human-like TMTFs.

### Differences among experimental conditions

Generally, the TMTFs in the lower layers were less similar to those in humans, and as the layer number went higher, the similarity also became higher (Figure 7, Figure 13b). More detailed inspections of individual stimulus parameters, however, indicate that changes in the form of TMTFs along the layers appeared to vary with the stimulus parameters. The change seems most prominent in the broadband carrier conditions, and least in the condition with the 314-Hz carrier bandwidth, showing almost constant TMTF forms across layers, followed by the 3- and 2-Hz bandwidth conditions (Figure 6, Figure 13a). In this study, we could not see a consistent relationship between the TMTF difference across layers and the stimulus parameters. There is a possibility that humans perform AM detection with different strategies in different experimental conditions. For example, the TMTF in the 314-Hz carrier bandwidth condition appeared similar to that in humans in as early as the first layer. This suggests that AM sensitivity in this condition might be explained only from the stimulus statistics. These results highlight the importance of testing a variety of stimulus parameters when investigating AM sensitivity in any system, including humans and computational models.

### Analyzing machine learning models with a combination of psychophysics and neurophysiology

From a machine learning point of view, this study can be viewed as an attempt to understand an NN with a combination of psychophysical and neurophysiological methods. Motivated by the fact that recent NNs have become too big and complicated for gaining an intuitive understanding of their mechanisms, a number of methods have been proposed for analyzing the NN’s behavior and stimulus representations (Montavon et al., 2018; Cammarata et al., 2020). The more difficult it is to understand an NN, the more important it would be to analyze it from a variety of perspectives. We can learn from a tradition of psychophysical and neurophysiological studies that have established various methods to investigate complicated biological systems (Eijkman, 1992; Leibo et al., 2018; Barrett et al., 2019; Richardwebster et al., 2019). The success of the present study demonstrates the utility of multidisciplinary analysis on a single platform.

## Methods

Code and results are available at https://github.com/cycentum/Human-like-Modulation-Sensitivity-through-Natural-Sound-Recognition.

### Model construction and evaluation

We used a multi-layer feed-forward NN as a model of the auditory system. The modeling framework was the same as in our previous study (Koumura et al., 2019). The architectural parameters (namely, the number of layers, number of units per layer, convolutional filter width, and convolutional dilation width) were newly sampled in this study. Each layer consisted of a dilated convolution (van den Oord et al., 2016) followed by an exponential linear unit (Clevert et al., 2016). Convolution was along the time axis. Above the topmost layer was a classification layer consisting of a convolution with a filter size of 1. In this way, the model worked as a fully-convolutional NN. The input time window was 0.2 s. In other words, the model estimated the sound category of every 0.2 s of the input sound. During optimization, softmax cross entropy for sound categories was computed at a single time step of the model output (corresponds to the input sampling rate) and the parameters (namely, convolutional weights and biases) were updated to minimize the error. During the evaluation, the output of the classification layer was averaged over time to estimate a single category per input sound segment.

Optimization was conducted with a standard backpropagation method using the Adam optimizer with a learning rate of 10^-4^. The sound data were divided into training and validation sets. We used the early-stopping strategy. This means that parameter update was conducted with part of the training set until recognition accuracy stopped improving for the other part of the training set.

#### Sound data for optimization

We used two datasets for optimization, ESC-50 (Piczak, 2015) and TIMIT (Garofolo et al., 1993). Both datasets are commonly used for sound recognition and are relatively small (Fonseca et al., 2022). We did not use larger datasets because our purpose was not to achieve state-of-the-art sound-recognition performance. The optimization to the two datasets and the analysis of the models were conducted independently.

ESC-50 defines 5 folds. We used folds 1 to 4 for training and fold 5 for validation. In the training, folds 1 to 3 were used for the parameter update and fold 4 was used for early stopping. Some sound clips end with absolute 0 amplitude values, probably to make the clip duration 5 s in otherwise shorter sounds. We excluded such 0 tailings.

TIMIT defines training and test sets. The test set includes sentences spoken by the core-test speakers and non-core-test speakers. For validation, we used sentences spoken by the core-test speakers. For training, we used the training set for the parameter update and the sentences spoken by the non-core-test speakers for early stopping. We excluded sentences included in both the training and test sets. This process ensured that there was no duplication of sentences or speakers in the training and validation sets.

Before being fed to the model, the sound signals were highpass filtered at 20 Hz, and the 10-ms raised-cosine ramps were applied to the onset and the offset. During training, the sound amplitude was slightly varied clip by clip. During the evaluation, it was fixed to the mean value of that for training.

#### Architecture search

For the architecture search, we tested architectures that varied in the number of layers, number of units per layer, and convolutional filter size and dilation width. The number of layers was 7, 9, 11, or 13. For each number, we sampled 20 models by varying the number of units per layer, convolutional filter size, and convolutional dilation width. The convolutional filter width and the dilation width were randomly sampled for each layer with the constraint on the input time window being 0.2 s. The number of units per layer was either 32, 64, 128, 256, or 512. To avoid an expensive computation of training all models over numerous iterations (Zhou et al., 2020), we conducted a two-step architecture search as follows. In the first step, we sought the number of layers that would potentially achieve the highest recognition accuracy. All models were trained until the recognition accuracy for a subset of the training set stopped improving for 32 epochs. Average recognition accuracy at this point of the four best models among those with the same number of layers was the highest for the 13-layer models (Figure 16). Thus, we selected the 13-layer architectures and discarded the others. In the second step, we further trained those 20 models until the recognition accuracy on a subset of the training set stopped improving for 96 epochs. We selected the four models with the highest recognition accuracy for the subsequent psychophysical and neurophysiological analyses.

**Figure 16.**
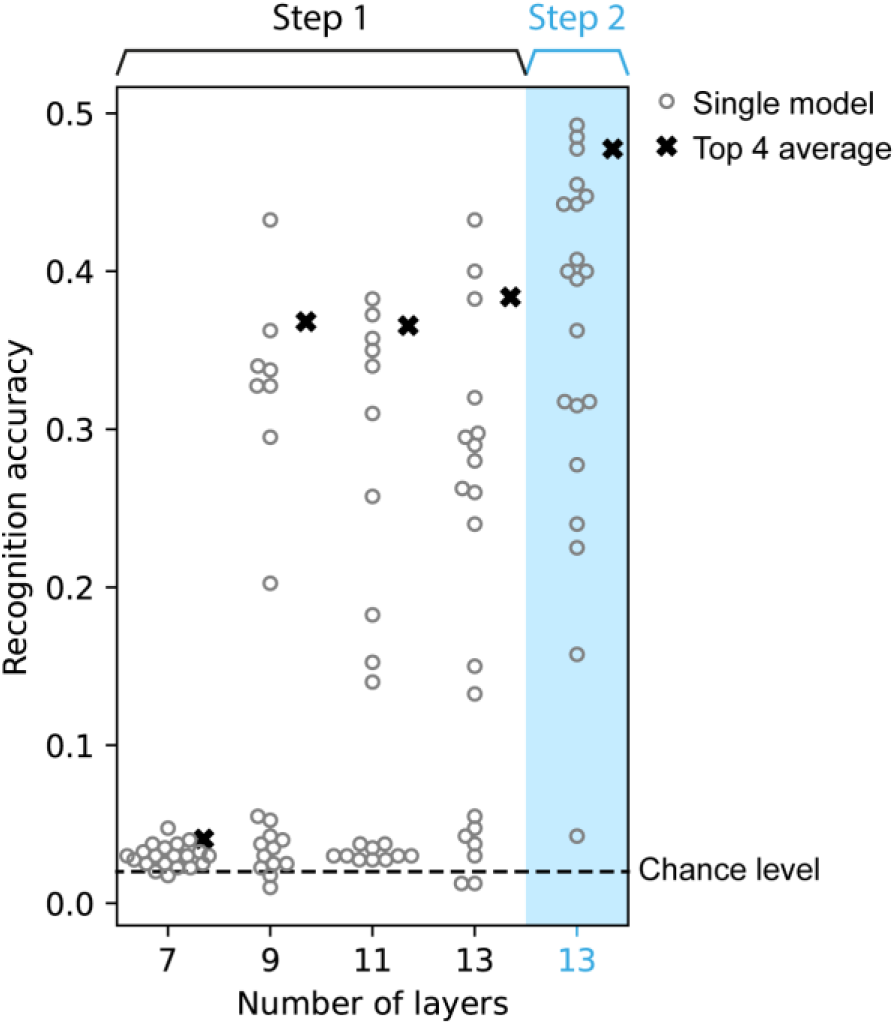
Sound recognition accuracy of the models with different architectures. In the first step of the search process, four models with 13 layers had the highest average accuracy (area with the white background). In the second step, the accuracy of the models with 13 layers improved after further optimization (area with the blue background).

### Psychophysics

To measure the AM detection threshold in the model, we simulated AM detection experiments in human psychophysics. To compare our results fairly with those produced by humans, the simulations duplicated the procedure of the human experiments as precisely as possible. One exception was that, in human studies, a detection threshold is estimated with a staircase method, whereas we computed the AM detection accuracy for each modulation depth independently.

We simulated a 2- or 3IFC task. In each trial, two or three stimuli were presented to the model, one of them being modulated, the others not. The task was to correctly identify the modulated interval. We conducted this task by assuming an AM detection process based on the model activities. We conducted 128 trials for each AM depth, from which the proportion of correct trials was calculated.

The proportion of correct trials plotted against the AM depth yields a psychometric curve (Figure 5c). It was fitted with an asymmetric sigmoid function (Richards, 1959; Fekedulegn et al., 1999). The detection threshold was defined as the AM depth at which detection accuracy was 70.7% on the fitted curve. In some conditions, the threshold could not be estimated because the proportion of correct responses was either too high or too low at all tested AM depths. Excluding such a condition would result in an overestimation of the similarity to human TMTFs. To avoid the overestimation, instead of excluding such a condition, the threshold was clipped to the maximum or minimum values of the tested range of the AM depth. The range is described below.

#### AM detection based on time-averaged unit activities

An xIFC task was conducted by estimating the modulated interval from model activities. Specifically, we assumed AM detection based on time-averaged unit activities (Figure 5a, b). For each stimulus interval, unit activities in the model were averaged over time. From the time-averaged activities in a single layer, a logistic regression was trained to estimate whether the stimulus was modulated. The proportion of correct trials was computed in a 4-fold cross validation of a total of 128 trials. In each of the 32 held-out trials, the probability of the stimulus being modulated was calculated for each stimulus interval, and the interval with the maximum probability was considered as the response in that trial. If that interval was actually the modulated interval, the trial was considered correct. L2 regularization was applied to logistic regression. The regularization coefficient was optimized in another 4-fold cross validation within the training set.

#### Stimulus

We tested six stimulus parameters from three independent human studies. All of them were sinusoidally amplitude modulated narrow- or broad-band white noise. The stimulus parameters and generation procedure were as closes as possible to those in the human studies, except for amplitude scaling: in the human studies, it was based on sound pressure levels, whereas in this study, it was based on the RMS. The stimulus RMS was adjusted to the average RMS of the training set. In the short-broadband condition, the stimulus amplitude was scaled before applying modulation (Viemeister, 1979). In the other conditions, it was scaled after modulation (Dau et al., 1997a; Lorenzi et al., 2001b, 2001a).

In the narrowband-carrier conditions, Gaussian noise was bandpass filtered with a digital Fourier transform. In the 314-Hz carrier bandwidth condition, bandpass filtering was applied after modulation (Dau et al., 1997a). In the other conditions, bandpass filtering was applied before modulation (Dau et al., 1997a; Lorenzi et al., 2001a). The Gaussian noise carrier was sampled independently in each stimulus.

AM depth is expressed in dB relative to the sound amplitude. In the case of sinusoidal AM, a sound with an AM depth *m* in dB and rate *f* is defined as,

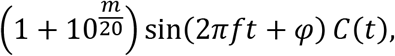

where *φ* is the AM starting phase, *t* is time, and *C*(*t*) is a carrier signal. The AM starting phase was fixed to 0 in the 3 Hz-, 31 Hz-, and 314 Hz-bandwidth conditions and in the long-broadband condition (Viemeister, 1979; Dau et al., 1997a). In the other conditions, it was randomly sampled independently in each stimulus (Lorenzi et al., 2001b, 2001a).

The range and steps of the AM rate differ among the human experiments. For each condition, we chose 8 AM rates evenly spaced on a log scale within the range in the particular human experiment. The AM depths ranged from −60 to 0 dB in the 2-Hz carrier bandwidth condition and −40 to 0 dB in the other conditions. They were spaced every 4 dB.

#### Quantitative comparison of model and human TMTFs

Previous human studies reported TMTFs in multiple human subjects. We took TMTF values from those studies and compared them with those in our model. Before calculating the quantitative similarities, we averaged the TMTFs across subjects. When the AM rates did not match among subjects, linear interpolation along the log-scaled AM rate was conducted.

Likewise, we averaged TMTFs of the four selected models. Then, we compared the averaged human TMTFs and averaged model TMTFs in terms of their relative patterns and absolute values. Similarity of their relative patterns was quantified by the pattern-similarity index, i.e., the correlation coefficient of the human and model TMTFs. Similarity of the absolute values was quantified by the discrepancy index, i.e., the RMS deviation:

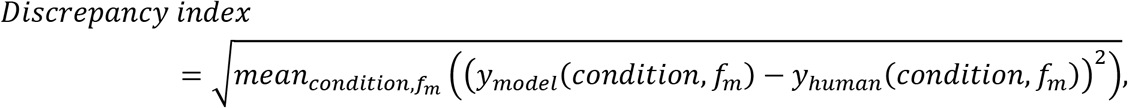

where *condition* and *fm* are the experimental conditions (size = 6) and the AM rates in each condition (size = 8), and *y* is a detection threshold.

The net difference between the human and model TMTFs was defined as the average signed difference between them.

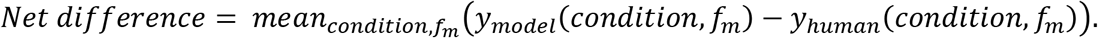

Positive/negative values of net difference mean larger/smaller thresholds in the model than in humans on average. To take all stimulus conditions into account, TMTFs in all conditions were pooled when calculating those indices.

#### Manipulation of the training data for exploring critical features

To evaluate the importance of Env and TFS, we made degraded versions of the training data by disrupting either the Env or TFS components of the sound.

Single-band Env signals were made by combining the Env component of a sound and a TFS component of white noise.

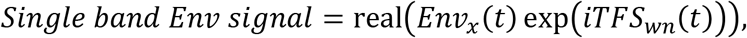

where *Env_s_* and *TFS_s_* are the Env and TFS components of a signal *s*, *x* and *wn* are the original sound and a white noise with the same RMS as *x*, *t* is time, *i* is the imaginary unit, and real converts a complex signal to its real part. The Env and TFS components are defined as the magnitude and phase of the Hilbert-transformed complex analytic signal.

Single-band TFS signals were made by flattening the Env component of a sound.

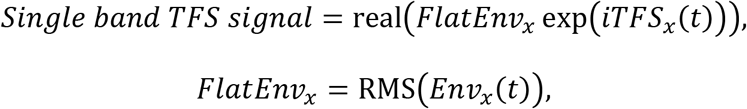

where *FlatEnv_s_* is a flattened Env of a signal *s*, which takes a constant RMS value of the Env component.

When making multi-band Env and TFS signals, we first decomposed the sound into subbands with a linear bandpass filterbank. The filter center frequencies ranged from 20 Hz to the Nyquist frequency and were spaced every 1 equivalent rectangular bandwidth (ERB) (Moore, 2012). Then, we computed the Env and TFS components for each subband. The Env or TFS components were disrupted in the same way as in the single-band signals, and the multiband signals were added together to form the final output:

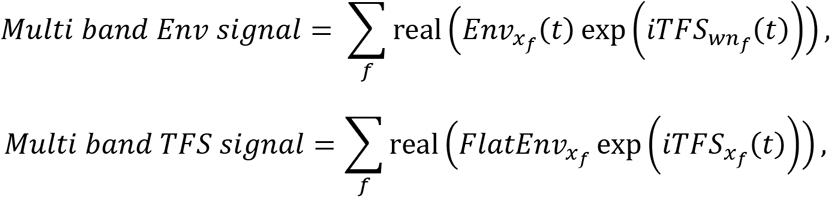

where *s_f_* is the *f^th^* subband of the frequency-decomposed signal *s*. Other than the difference in the training data, the procedures of the optimization and analysis were completely the same as the models trained on the original sounds.

#### AM detection based on template correlation

For each sound interval in the xIFC task, the correlation was calculated between the unit activities in a layer in response to the stimulus and the template. The interval with the largest correlation was taken to be the response to the trial. The correlation was defined by the sum of products as in the previous study (Dau et al., 1997a). It was calculated for all units in each layer:

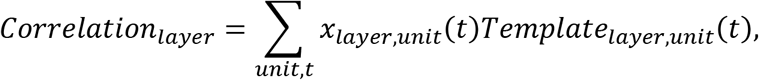

where *x_layer,unit_* is the activity in a specific unit in the target layer, and *t* is time.

To make a template, first, we averaged the unit activities across 128 independent fully-modulated and non-modulated stimuli. The template was defined as average unit activities for fully-modulated stimuli minus the average unit activities for non-modulated stimuli (Dau et al., 1997a):

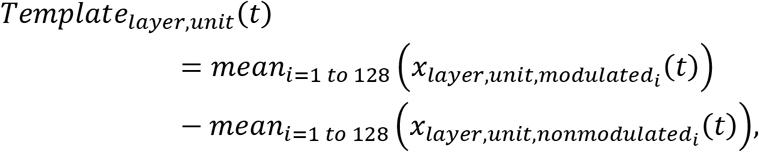

where *X_layer,unit,modulatedi_* and *x_layer,unit,nonmodulatedi_* are the unit activities in response to the *i^th^* modulated and non-modulated stimulus, respectively.

### Neurophysiological similarity between NN layers and brain regions

The neurophysiological similarity between NN layers and brain regions was computed in the same way as in our previous study, except that the resolution of the AM rates at which AM tuning was computed was decreased in this study for reducing the computational cost. A detailed description of the method is provided in our previous paper (Koumura et al., 2019).

Unit activities in the model were recorded while presenting it with sinusoidally amplitude-modulated broadband white noise. The AM tuning was defined in terms of the time-averaged unit activities and the synchrony of the activities to the stimulus modulation. It was characterized by the best AM rate and the upper cutoff rate. The best AM rate was defined as the AM rate at which the tuning curve reached a maximum. The upper cutoff rate was defined as the AM rate at which the tuning started to decrease. Distributions of best and upper cutoff rates were compared between NN layers and brain regions. Similarity between an NN layer and a brain region was defined as 1 minus the Kolmogorov–Smirnov distance of the distributions. The AM tuning in the ANS was taken from previous neurophysiological studies (Müller-Preuss, 1986; Langner and Schreiner, 1988; Schreiner and Urbas, 1988; Batra et al., 1989; Preuß and Müller-Preuss, 1990; Frisina et al., 1990; Joris and Yin, 1998, 1992; Rhode and Greenberg, 1994; Zhao and Liang, 1995; Bieser and Müller-Preuss, 1996; Condon et al., 1996; Schulze and Langner, 1997; Eggermont, 1998; Huffman et al., 1998; Joris and Smith, 1998; Kuwada and Batra, 1999; Krishna and Semple, 2000; Lu and Wang, 2000; Lu et al., 2001; Liang et al., 2002; Batra, 2006; Zhang and Kelly, 2006; Bartlett and Wang, 2007; Scott et al., 2011; Yin et al., 2011).

## Acknowledgments

This work was supported by JSPS KAKENHI Grant Number JP20H05957 (Grant-in-Aid for Transformative Research Areas (A) “Analysis and synthesis of deep SHITSUKAN information in the real world”).

